# Performance of computational algorithms to deconvolve heterogeneous bulk tumor tissue depends on experimental factors

**DOI:** 10.1101/2022.12.04.519045

**Authors:** Ariel A. Hippen, Dalia K. Omran, Lukas M. Weber, Euihye Jung, Ronny Drapkin, Jennifer A. Doherty, Stephanie C. Hicks, Casey S. Greene

## Abstract

**Background:** Single-cell gene expression profiling provides unique opportunities to understand tumor heterogeneity and the tumor microenvironment. Because of cost and feasibility, profiling bulk tumors remains the primary population-scale analytical strategy. Many algorithms can deconvolve these tumors using single-cell profiles to infer their composition. While experimental choices do not change the true underlying composition of the tumor, they can affect the measurements produced by the assay.

**Results:** We generated a dataset of high-grade serous ovarian tumors with paired expression profiles from using multiple strategies to examine the extent to which experimental factors impact the results of downstream tumor deconvolution methods. We find that pooling samples for single-cell sequencing and subsequent demultiplexing has a minimal effect. We identify dissociation-induced differences that affect cell composition, leading to changes that may compromise the assumptions underlying some deconvolution algorithms. We also observe differences across mRNA enrichment methods that introduce additional discrepancies between the two data types. We also find that experimental factors change cell composition estimates and that the impact differs by method.

**Conclusions:** Previous benchmarks of deconvolution methods have largely ignored experimental factors. We find that methods vary in their robustness to experimental factors. We provide recommendations for methods developers seeking to produce the next generation of deconvolution approaches and for scientists designing experiments using deconvolution to study tumor heterogeneity.

## Background

Solid tumors are highly heterogeneous tissues; the malignant cancer cells cohabitate and interact with various immune and stromal cells, known broadly as the tumor microenvironment (TME), in complex ways [1]. For cancer patients with the same tumor type, differences in the TME can yield different outcomes in progression, treatment response, and overall survival. TME composition affects immune cells’ ability to locate and kill malignant cells, the bioavailability and effectiveness of chemotherapy drugs, the availability of oxygen and other nutrients needed for cancer cell growth, and the possibility of metastasis [2, 3]. For these reasons, thorough characterization of the TME is an active area of cancer research [4, 5].

Researchers often use Bulk RNA sequencing (RNA-seq) and single-cell RNA-seq (scRNA-seq) to examine the TME. Bulk sequencing—extracting RNA from pulverized tissue—is cost-effective and allows for transcriptome-wide coverage of total RNA. Many large cancer characterization efforts, such as The Cancer Genome Atlas, have bulk RNA-sequenced hundreds or thousands of samples [6]. Unfortunately, bulk RNA-seq loses direct information on tumor purity and cell type composition. Single-cell RNA-seq involves dissociating tissue and characterizing individual cells, retaining cell type-specific information. However, scRNA-seq is expensive and thus hard to scale to large datasets. scRNA-seq also produces much sparser data than bulk RNA-seq [7]. Each data modality presents unique experimental opportunities and challenges, but it is possible to combine bulk and single-cell data to computationally estimate tissue composition of bulk RNA-seq data using single-cell profiles, providing estimates of the TME for larger studies.

In the context of the TME, deconvolution describes the challenge of estimating cell type abundances from bulk profiles. Methods can be reference-free [8–10] or reference-based [11–15]. Many reference-based methods use a matrix of signature marker genes, but with the advent of single-cell sequencing, reference-based methods using profiles drawn from single-cell observations have become widespread []. We focus on reference-based methods in this paper.

Whether or not methods use single-cell data as input, many within-method validations and cross-method benchmarks rely on single-cell data to assess the accuracy of a deconvolution method [16–18]. These assessments aggregate scRNA-seq data to create simulated or “pseudo-bulk” tumors with known cell type proportions. This assumes that single-cell and aggregated bulk data are biologically equivalent and that performing well on one data type indicates capturing similar information on the other. However, there are several technical differences that strain this assumption.

One source of technical variability between single-cell and bulk sequencing is dissociation. Separating cells from each other requires vigorous chemical and/or physical digestion, which can lyse cell membranes or otherwise compromise cell integrity [19]. Certain cell types are more sensitive to this process and are systematically underrepresented in scRNA-seq data [20]. Deconvolution algorithms that assume complete representation of cell types may perform well on pseudo-bulk assessments but could underperform in practice.

Another difference between single-cell and bulk RNA sequencing is the method of mRNA enrichment. Most RNA in any given cell is ribosomal RNA, which is undesired in most RNA-seq studies [21]. There are two prevailing ways to enrich for non-ribosomal RNA [22]. Many bulk RNA-seq experiments use ribosomal depletion which directly removes rRNA from a sample. This approach performs well for capturing partially-degraded RNA, such as that found in formalin-fixed paraffin-embedded (FFPE) tissue [23]. An alternative strategy is poly-A capture, which adds primers that ligate to the polyadenylated 3’ ends of mRNA. Many single-cell protocols use poly-A-based methods. It is unknown how using reference profiles from poly-A captured single cells affects the deconvolution of rRNA-depleted bulk samples.

In this work, we generate a unique dataset of high-grade serous ovarian tumors and use it to directly examine the effects of protocol differences and their ramifications for deconvolution. We focus specifically on tumor deconvolution. While some deconvolution methods are designed for cancer data [11, 17], benchmarks have been performed predominantly on normal tissue [16, 18]. Solid tumors present unique challenges in deconvolution. Aberrant and dysregulated tissue growth often yields incomplete dissociation with many cells damaged [24]. Inter-patient heterogeneity is also much greater for malignant cells than for normal cell types [25], making it harder to generalize patterns across samples. Indeed, robustness to the noise contributed by the tumor fraction has been called one of the major challenges deconvolution algorithms face [26]. We compare the consistency of six deconvolution methods across protocols and assess their accuracy on cancer data. Finally, we propose a series of recommendations for researchers looking to sequence cancer samples for use in deconvolution and subsequent at-scale studies of the TME.

## Results

### Experimental design

Our dataset comprises tumor data from n=8 high-grade serous ovarian carcinoma (HGSOC) patients. HGSOC is known to have considerable inter-patient and intra-tumor heterogeneity, making deconvolution particularly valuable [27–29]. In addition, HGSOC tumors exemplify the kinds of challenges faced in cancer sequencing. Since HGSOC easily disseminates through the peritoneal cavity and forms small metastases, most debulking surgeries are extensive and take many hours [30], increasing the RNA and tissue degradation prior to freezing or fixture. The tumor’s histopathology is marked by extensive regions of necrotic tissue [31] resulting in a large amount of cellular debris at sequencing. Also, HGSOC cells have high genomic instability and a particularly high burden of copy number variants [32, 33] which can complicate deconvolution. By focusing on a challenging tumor type, we aim to identify best practices that are robust to real-world experimental conditions and thus have relevance to many other solid tumor types.

We used data from eight HGSOC tumors for which frozen tumor chunks and frozen dissociated cells were available (Methods). To directly assess the ways different library preparation methods affect deconvolution in cancer data, we assayed our data in multiple ways (Figure 1). We performed RNA extraction on the tumor chunks, enriched for mRNA with rRNA depletion, and performed bulk RNA-sequencing. We will refer to this data type as “rRNA^-^ Chunk.” Ribo-depletion on undigested tissue is one of the most common protocols for cancer RNA-seq datasets and is thus likely to be used as an input for deconvolution. We also performed rRNA depletion on dissociated cells and performed bulk RNA-sequencing. We will refer to this data type as “rRNA^-^ Dissociated.” By comparing the rRNA^-^ Chunk and rRNA^-^ Dissociated data, we examine the effect of dissociation without other confounding factors that would be involved in bulk vs. single-cell comparisons. We also performed poly-A (3’) capture and performed bulk RNA-sequencing on RNA from dissociated cells. We will call this data type “polyA^+^ Dissociated.”

**Figure 1:**
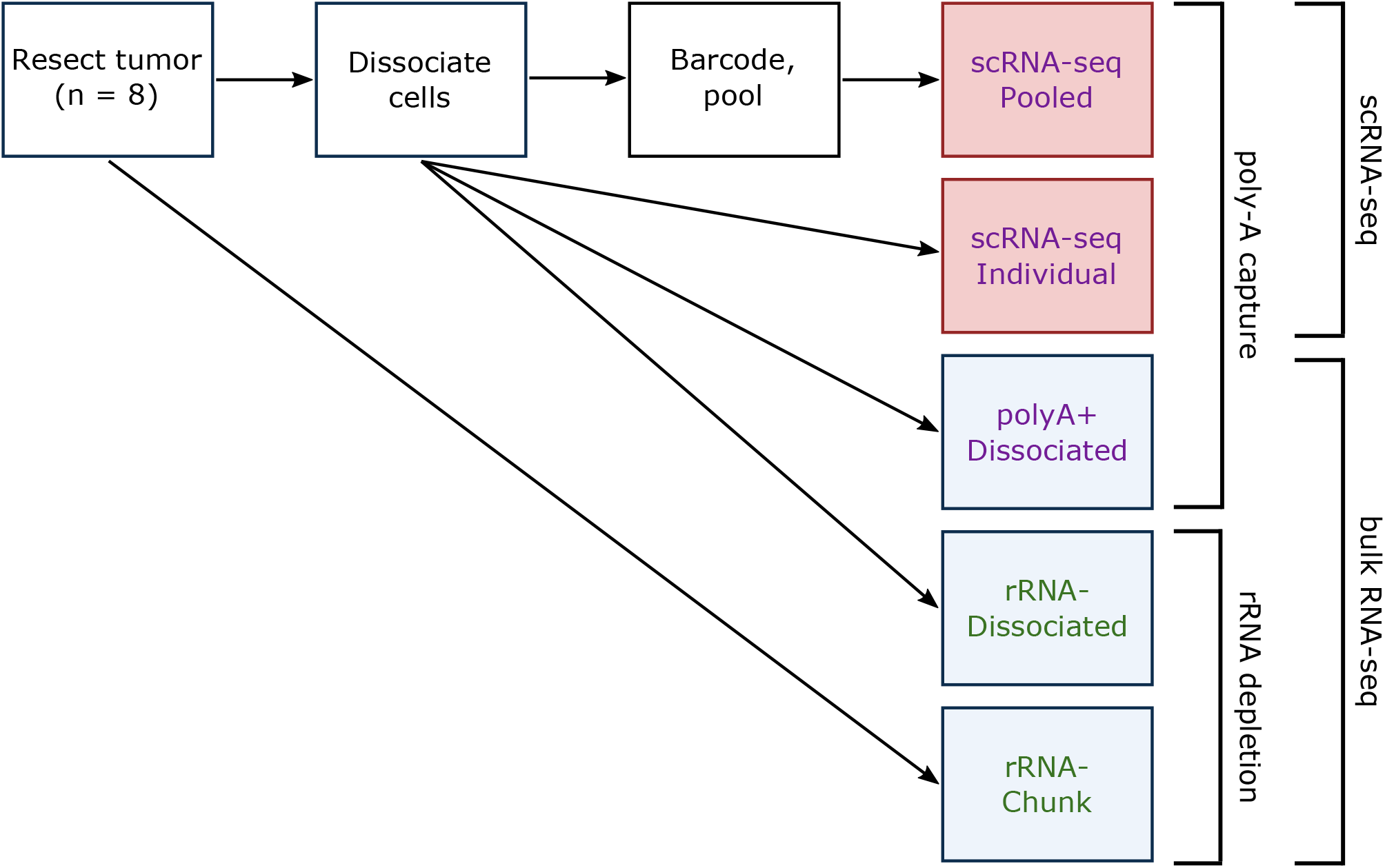
Overview of experimental design. Each tumor was profiled in five different ways, three times with bulk RNA-seq and twice with scRNA-seq using two strategies for mRNA enrichment, rRNA depletion and poly-A capture.

In addition to our three bulk sequencing data types, we performed two different scRNA-seq assay types. For one portion of the dissociated cells, we performed scRNA-seq on each tumor separately. We will refer to these as the “scRNA-seq Individual” samples. For another portion of the dissociated cells, we added a barcoded antibody and pooled the cells into batches (two sets of four samples each), and performed scRNA-seq on the pools. We will refer to these as the “scRNA-seq Pooled” samples. Performing scRNA-seq both individually and in pools allows us to directly compare deconvolution results using reference profiles from each data type and also evaluate the impact of demultiplexing on deconvolution.

### Multiplexing increases throughput and preserves sample-specific information

Pooling has the potential to greatly increase the scalability of single-cell profiling, but it introduces technical and computational challenges. Pooled samples require a higher total number of cells to be loaded for acceptable coverage of each sample. In cancer samples with high cellular debris from necrotic tissue, loading more cells may increase the risk of clogging the microfluidic device. By adding an extra debris-filtering step on the batched samples (Methods), we could sequence cell counts comparable to or higher than the individual single-cell runs (Table 1).

**Table 1:** Single-cell count per sample. All numbers are after filtering based on percentage of mitochondrial reads.

| Sample ID | Sample Type | Cells | Component samples |
| --- | --- | --- | --- |
| 2251 | Individual | 9464 |  |
| 2267 | Individual | 5345 |  |
| 2283 | Individual | 7627 |  |
| 2293 | Individual | 10609 |  |
| 2380 | Individual | 6300 |  |
| 2428 | Individual | 283 |  |
| 2467 | Individual | 6729 |  |
| 2497 | Individual | 9313 |  |
| A | Pooled | 7358 | 2267, 2283, 2293, 2380 |
| B | Pooled | 9814 | 2251, 2428, 2467, 2497 |

Upon successful sequencing, another challenge arises: identifying from which sample each cell originates. The process of computationally splitting the cells into groups by sample or patient of origin is known as demultiplexing. To determine if it’s possible to sufficiently demultiplex cells from cancer tissue to use them as reference profiles for deconvolution, we performed two kinds of demultiplexing: hash demultiplexing and genetic demultiplexing.

#### Hash demultiplexing is precise but limited at default thresholds

For hash demultiplexing, cells are labeled with an antibody targeting ubiquitous cell surface epitopes attached to a unique oligo-tag (one for each sample) and are then pooled; after sequencing, the tag on each cell is used to recapitulate the sample of origin [34]. We used 10X Genomics’ cellranger multi platform to do this. When performing demultiplexing based on antibody hashing in the two batches, 4246 and 3734 of the cells respectively (57.7% and 38.0%) were assigned to one sample, with 286 and 145 (3.9% and 1.5%) cells called as multiplets and 2823 and 5935 (38.4% and 60.5%) cells unassigned (Figure 2A-B, Table S1-S2).

**Figure 2:**
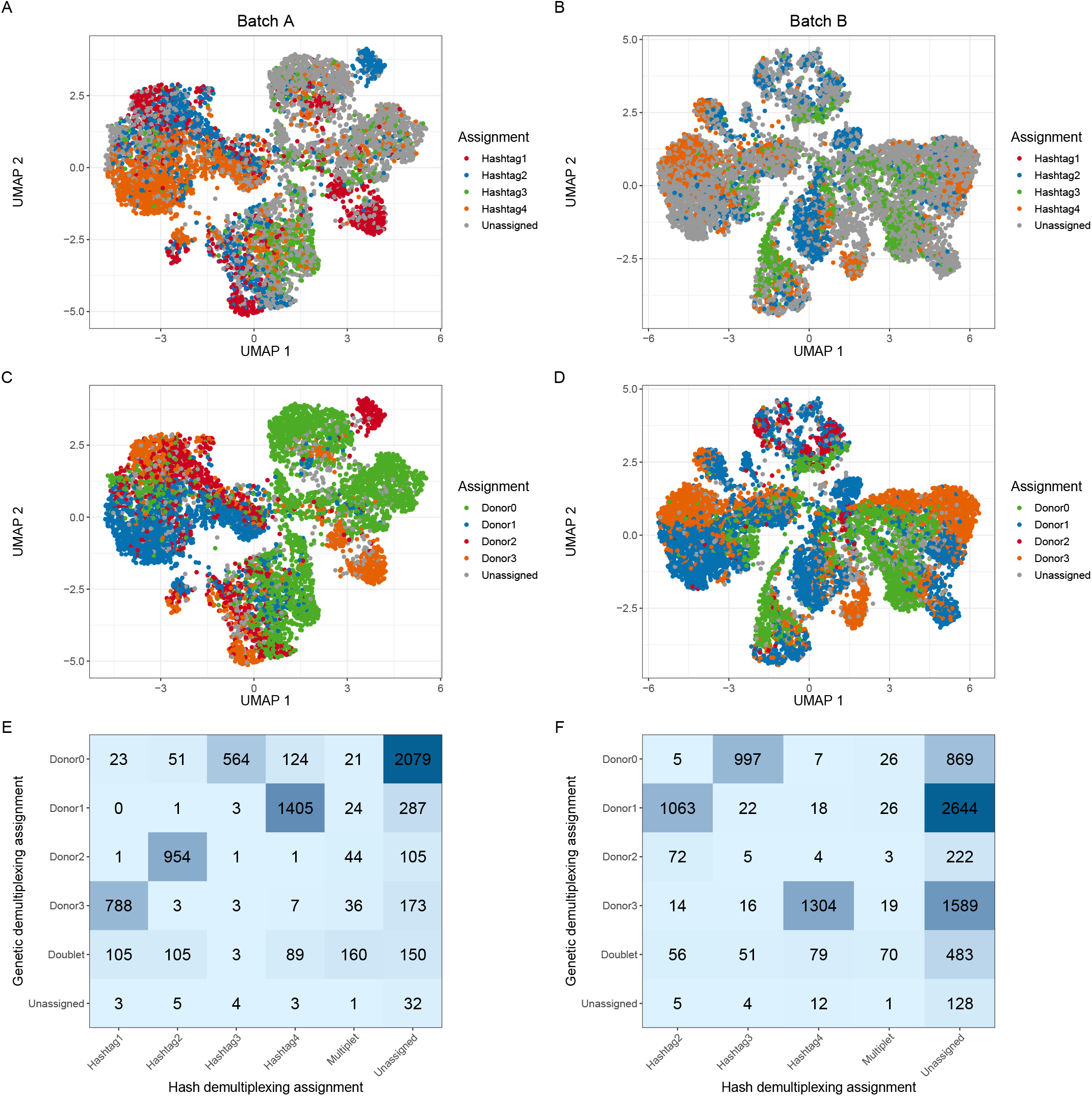
Results of antibody-based and genetic demultiplexing are concordant in cancer data. A-B) A UMAP representation of the pooled data from Batch A (A) and Batch B (B), colored by antibody-based assignment from cellranger multi. C-D) The same samples colored based on genetic demultiplexing assignment from vireo. E-F) A confusion matrix showing the overlap of assignments with antibody-based and genetic demultiplexing in each sample.

When reviewing the assignment probabilities for each cell, we found that many unassigned cells mapped to one antibody hashtag with reasonably high probability. The default assignment threshold for cellranger multi is 90% probability or greater of originating from one sample. When we relaxed this threshold to 85% probability, 4974 and 3831 cells were assigned (67.6% and 39.0% of total) (Figure S1A-B). A further relaxed threshold of 80% probability yielded 5536 and 4032 assigned cells (75.2% and 41.1% of total) (Figure S1C-D).

Given the high number of unassigned cells under default parameters, we checked if there was differential antibody adhesion based on cell type, which if present could bias downstream deconvolution. We assigned a cell type label to all pooled cells using the CellTypist package [35] combined with unsupervised clustering (Methods). We found across tested probability thresholds that epithelial cells and fibroblasts were proportionally more likely to be unassigned in Batch A, whereas T cells were proportionally less likely to be unassigned (Figure S2A). We did not observe a similar bias in Batch B (Figure S2B). One difference between these two batches is that most of the unassigned cells in Batch A were assigned to a single sample (id 2283) when the probability threshold was relaxed. In contrast, the newly-assigned cells at lower thresholds were more evenly distributed in Batch B, suggesting lower overall antibody adhesion in the cells from sample 2283. We posit that in samples where overall antibody adhesion is low, perhaps due to insufficient reagent or insufficient time for adhesion, antibodies are preferentially likely to bind to the cell surface markers of certain cell types, perhaps through greater prevalence or steric availability of CD298 and/or *β*2 microglobulin. Given sufficient time or reagent, however, we posit that the antibodies will eventually bind to all cell types, explaining the lack of cell type bias in other samples/batches. This emphasizes the importance of titrating reagents based on the amount of cellular input, as recommended in the cell multiplexing procol we used [36]. These results also highlight the need to scrutinize the data post-sequencing and test a range of assignment thresholds rather than simply relying on default parameters to maximize the number of confidently assigned cells.

#### Genetic demultiplexing performs well in HGSOC samples

We also performed genetic demultiplexing of the pooled cells. Instead of identifying sample of origin based on an experimentally-added antibody, genetic demultiplexing leverages unrelated patients’ innate genetic variation to group cells based on their genotype [37]. Genotypes can be called using common variants from publicly available data, e.g., from the 1000 Genomes Project, or with genotypes called from another data modality in the same samples. The latter allows cells to be directly mapped back to samples rather than arbitrarily labeled. We used bcftools to genotype our bulk RNA-seq data [38], cellSNP-lite to genotype the single cells [39], and vireo to use the called genotypes to assign a sample of origin [40]. Under this framework, we assigned 6730 and 8866 of the cells respectively (91.4% and 90.3%) to one sample, with 558 and 705 (7.6% and 7.2%) called as multiplets and 70 and 243 (1.0% and 2.5%) unassigned (Figure 2C-D).

Since genetic demultiplexing relies on the ability to call sample-specific genotypes for common variants within single cells, the inherent genomic instability of cancer cells has previously been an area of concern. Simulated experiments have indicated that genetic demultiplexing was possible in tumor samples [41], and these results offer confirmation in real experimental data. It has been shown that using genotypes from bulk data from the sample samples (when available) is preferable for cancer demultiplexing [41]. One could imagine that the selection strategy used for bulk data could affect results—for example, by unevenly sampling across transcripts. Here we found that genotypes from paired bulk RNA-seq samples appear to be highly consistent across protocol types. We performed genotyping on our three bulk RNA-seq datasets (rRNA^-^ Chunk, rRNA^-^ Dissociated, polyA^+^ Dissociated) and performed genetic demultiplexing with each as a reference. We found that over 99% of cells had the same genetic demultiplexing assignment in each run (Figure S3A-C).

#### High concordance between hash and genetic demultiplexing

Encouragingly, we saw a high degree of overlap in assignments between hash and genetic demultiplexing. In cells assigned to a sample by both methods, 94.3% and 95.2% of cells were assigned to the same sample (Figure 2E-F). The biggest area of discordance overall was that many cells were assigned by genetic demultiplexing and left unassigned by hash demultiplexing. This effect was somewhat lessened at the more permissive 80% cellranger multi threshold. We attribute the higher number of assigned cells with genetic multiplexing, even after the relaxed hash assignment threshold, to incomplete adhesion of the antibody tags. For this reason, we elected to use cells assigned by genetic demultiplexing as our single-cell reference profiles for our main deconvolution analyses.

### Dissociation causes loss of certain cell types

As mentioned previously, some cell types are more resilient to dissociation than others, which can create a bias in deconvolution when using single cells as reference profiles and in comparison of bulk to single-cell data. We assessed the effect of dissociation on tumor transcriptomic data by comparing two of our bulk RNA-seq datasets: rRNA^-^ Chunk and rRNA^-^ Dissociated.

Principal component analysis (PCA) of the samples’ expression profiles revealed that individual sample types tended to segregate together in the first two principal components rather than based on dissociation status (Figure 3A). This indicated that inter-patient heterogeneity is still strongly present before and after dissociation. We ran differential expression using the DESeq2 package [42] (Figure 3B). The genes with the highest log fold-change of expression in the tumor chunks compared to the dissociated cells were hemoglobin genes (HBA1, HBA2, HBB). Hemoglobin genes were significantly reduced across all rRNA^-^ Dissociated samples when compared to their rRNA^-^ Chunk counterparts (Figure 3C). Erythrocytes (red blood cells) are the predominant expressors of hemoglobin, and are lysed and removed by many dissociation protocols [43], including the one that we used. We plotted other erythrocyte-specific genes [44] and found several were significantly more abundant in the tumor chunks as well.

**Figure 3:**
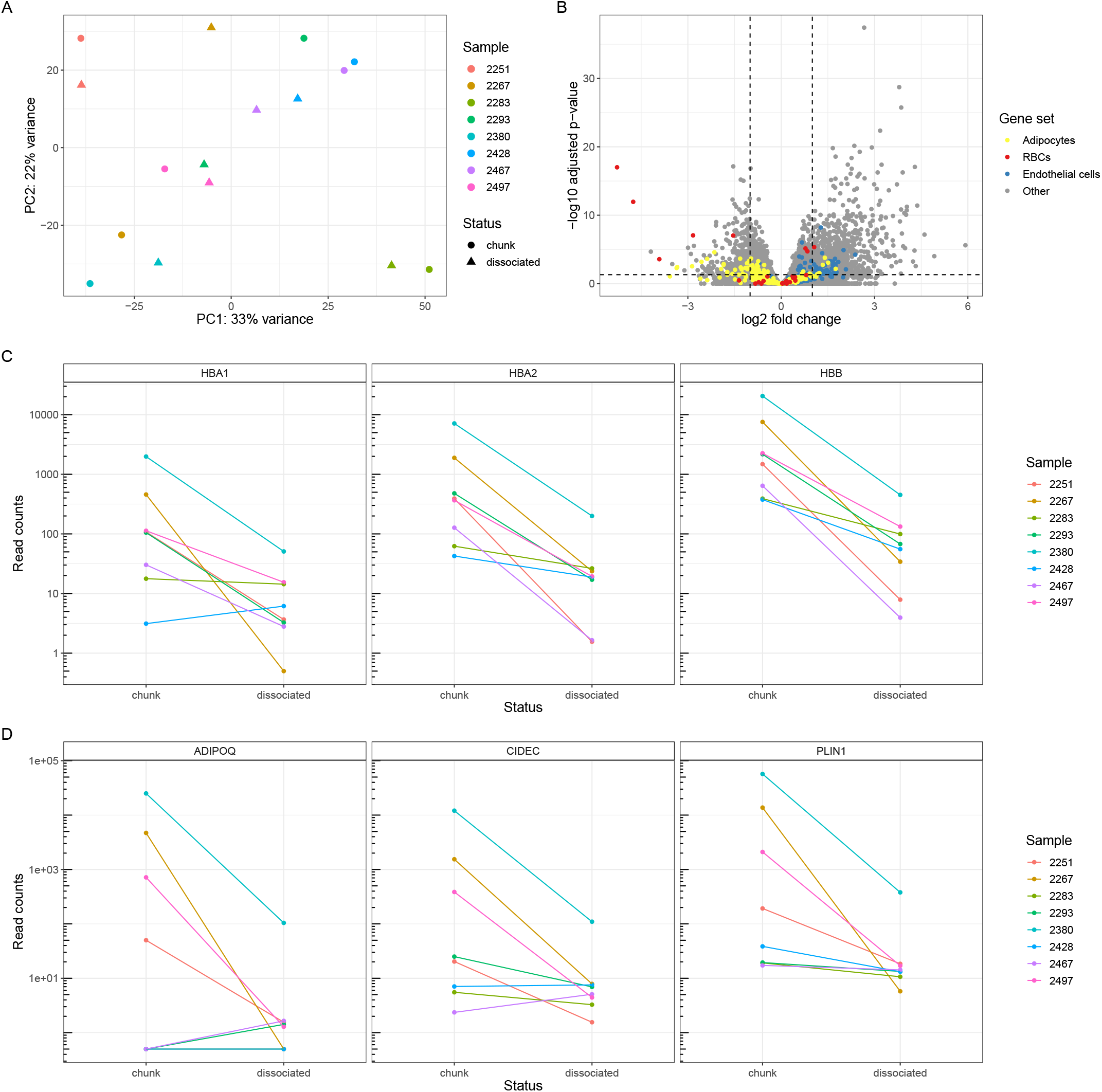
Dissociation causes loss of certain cell types. A) A principal component analysis of the rRNA^-^ Chunk and rRNA^-^ Dissociated bulk samples, where color indicates patient of origin and shape indicates dissociation status. Two points that are closer together on the PCA plot are more similar in their expression profiles. B) A volcano plot of the differential expression results based on dissociation status, with gene sets of interest colored: genes known to be upregulated in adipocytes [46], endothelial cells [46], and red blood cells [44]. C) Expression of hemoglobin genes in each sample based on dissociation status. D) Expression of selected adipocyte-related genes based on dissociation status.

Several other highly-increased genes in the tumor chunks were associated with adipose tissue (Figure 3B). Adipocytes are fragile and rarely survive dissociation [45]. In a comparison of single-cell and single-nucleus RNA-seq data, adipocytes were abundant in single-nucleus data and essentially absent from single-cell data [46]. Some of the samples expressed adipose-related genes in the tumor chunks but less in the dissociated cells. In other samples, adipose gene expression was low in both tumor chunks and dissociated cells (Figure 3D). These data support a model where some tumors have high numbers of adipocytes, which are lost during dissociation, and others lack substantial adipose tissue. While the surgical excision site was not recorded for these samples, our data are consistent with certain samples being derived from the omentum (a layer of fat lining the peritoneal cavity to which ovarian cancer cells preferentially migrate and colonize [47]) and others from other sites.

The omentum is a layer of fat lining the peritoneal cavity, to which ovarian cancer cells preferentially migrate and colonize [47]. While surgical excision site was not recorded for our samples, we suspect that the tissue samples with high adipocyte gene expression in the tumor chunks derive from the omentum, while the samples with little adipocyte gene expression derive from other sites.

Many genes were more abundant in the dissociated cells compared to the tumor chunks. These genes are related to a variety of biological pathways and cell types. Gene set expression analysis using cell type signature genes from the Molecular Signatures Database [48] showed that endothelial cells, fibroblasts, macrophages, and other immune cell types are more abundant in dissociated cells (Figure S4). We confirmed increased endothelial expression using marker genes from Emont et al [46] (Figure 3B). Indeed, many of the stromal and immune cell types one would expect to see in an HGSOC tumor are more abundant in the dissociated cells. We hypothesize this is due to increased relative abundance rather than a true biological enrichment. When red blood cells and adipocytes are removed, the relative abundance of markers of the remaining cell types necessarily increases.

While some cell type bias in dissociated cells is caused by easily avoided technical artifacts—one could alter their dissociation protocol to not include a red blood cell lysis step—others are not easily remedied, such as the loss of adipocytes and other fragile cell types. Other cancer types may also have other relevant cell types that are lost in dissociation, such as mesothelial cells [46]. Deconvolution using single-cell data for reference profiles will, at best, be unable to detect the presence of or quantify these fragile cell types. This poses a particular problem for adipocytes in ovarian cancer studies, where adipocytes are posited to have a direct role on tumor growth and metastasis [49, 50] and explain some aspects of inter-patient heterogeneity. Methods that assume all cell types have a reference present may exhibit unstable performance as they minimize residuals that arise from the absent cell types.

### mRNA enrichment affects gene abundance

Many deconvolution experiments use bulk data that has been ribosomal RNA-depleted and single-cell reference profiles that have been poly-A captured. While both poly-A capture and rRNA depletion have been shown to effectively enrich for mRNA across a variety of contexts [23, 51], it is not known if this experimental difference has a downstream effect on deconvolution. Comparing two of our datasets, rRNA^-^ Dissociated and polyA^+^ Dissociated, allows us to observe the impact different mRNA enrichment methods have on gene expression profiling.

To visualize the differences across samples and across data types, we used PCA on a regularized log-transformed dataset comprising all genes in the rRNA^-^ Dissociated and polyA^+^ Dissociated samples. We found that the first principal component segregated samples by patient, while second principal component completely separated the rRNA^-^ Dissociated and polyA^+^ Dissociated samples from each other (Figure 4A). The choice of mRNA enrichment method exerts a substantial effect on overall gene expression.

**Figure 4:**
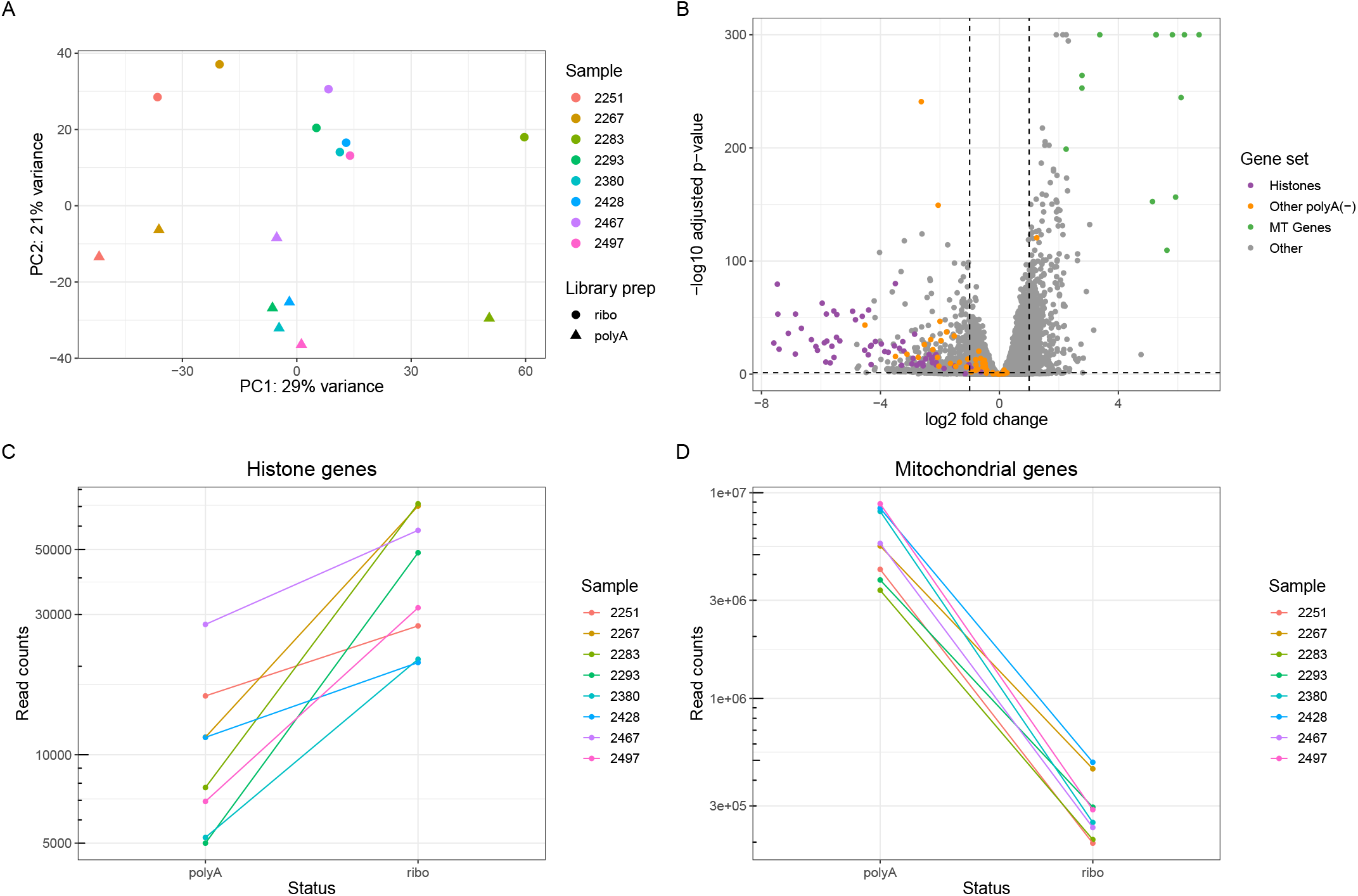
Method of mRNA enrichment affects gene expression profile. A) A principal component analysis of the rRNA^-^ Dissociated and polyA^+^ Dissociated bulk samples, where color indicates patient of origin and shape indicates whether samples were poly-A captured or rRNA depleted. Two points that are close together on the PCA plot are similar in their expression profiles. B) A volcano plot of the differential expression results based on mRNA enrichment method, with gene sets of interest colored: histones, other non-polyadenylated genes [55], and mitochondrial genes. C) Combined expression of all histone genes in each sample based on mRNA enrichment method. D) Combined expression of all mtDNA genes based on mRNA enrichment method.

We performed differential expression analysis to identify trends in global expression profiles (Figure 4B). Of the top 20 most differentially abundant genes (based on log fold change) in the ribo-depleted samples, 10 encoded histone proteins. To see if this effect was widespread among all histone genes, we aggregated their counts and found 1.7-fold to 10-fold enrichment of histone genes in the rRNA^-^ Dissociated samples compared to the polyA^+^ Dissociated samples from the same tumor (Figure 4C). There is a simple explanation for this: canonical histone RNAs are not polyadenylated and thus missed by poly-A capture protocols [52]. (The histone reads we observe in the polyA^+^ Dissociated samples are likely attributable to variant histones that are not cell cycle dependent and are polyadenylated.) Several other non-polyadenylated transcripts, such as TERC (the RNA component of telomerase) and RMRP (an endoribonuclease implicated in cancer progression [53, 54]), were also in the top 20 most differentially abundant genes in the rRNA^-^ Dissociated samples [55]. While these genes are not documented marker genes for cell types, researchers should expect that these genes will be substantially undercounted or missing in poly-A captured samples, which includes many existing tumor maps.

We observed another trend in the opposite direction: of the top 20 most differentially abundant genes in the poly-A captured samples, 10 originated from the mitochondrial transcriptome (mtRNA). Examining the aggregated counts of all mitochondrial RNA in both sets of samples, we found a 10-fold to 30-fold increase in mtRNA reads in poly-A samples compared to their ribo-depleted counterparts (Figure 4D). We initially found these results surprising since most mitochondrial RNAs have no connection to ribosomal machinery. However, the widely-used kit that we used for rRNA depletion has an off-target effect where non-ribosomal mitochondrial transcripts are depleted along with the mitochondrial ribosomes [56]. The apparent increased abundance of mitochondrial genes in poly-A samples is likely attributable to this technical artifact.

The percent of mtRNA reads is a major metric used for quality control of scRNA-seq data. Dissociation can result in a rupture of the cell membrane and loss of cytoplasmic RNA, causing an increase in the proportion of mitochondrial RNA [57]. Cells above a certain mitochondrial threshold are usually removed from analysis, assumed to be dead or irreparably compromised. If a researcher uses paired bulk RNA-seq that has been ribo-depleted as a reference for the expected fraction of mtRNA reads, they may choose an overly conservative threshold and lose many potentially informative cells.

### Assessing deconvolution accuracy and robustness together improves method evaluation

With more information on the effect different experimental decisions have on the data directly, we assessed the extent to which those experimental factors affect tumor deconvolution. We applied several commonly used deconvolution methods to our tumor data (Table 2) [11–15, 58]. We chose methods that return proportions of cell types, allowing us to directly compare results across methods. Each method has its own particular required inputs. In the case of methods that do not use scRNA-seq data, we used the marker gene matrices provided by the respective methods [14]. Most methods we examined require single cells as cell type reference profiles [11–13, 15, 58]. For these methods, we used cells from our pooled single-cell data that could be assigned to a sample by genetic demultiplexing and confidently annotated to a cell type (n=14,608). Having two types of single-cell data from the same samples (scRNA-seq Pooled and scRNA-seq Individual) allowed us to provide the pooled cells as a reference profile and leave the individually-sequenced cells to be used for validation without the pitfalls of using the same data for reference profiles and assessment.

**Table 2:**
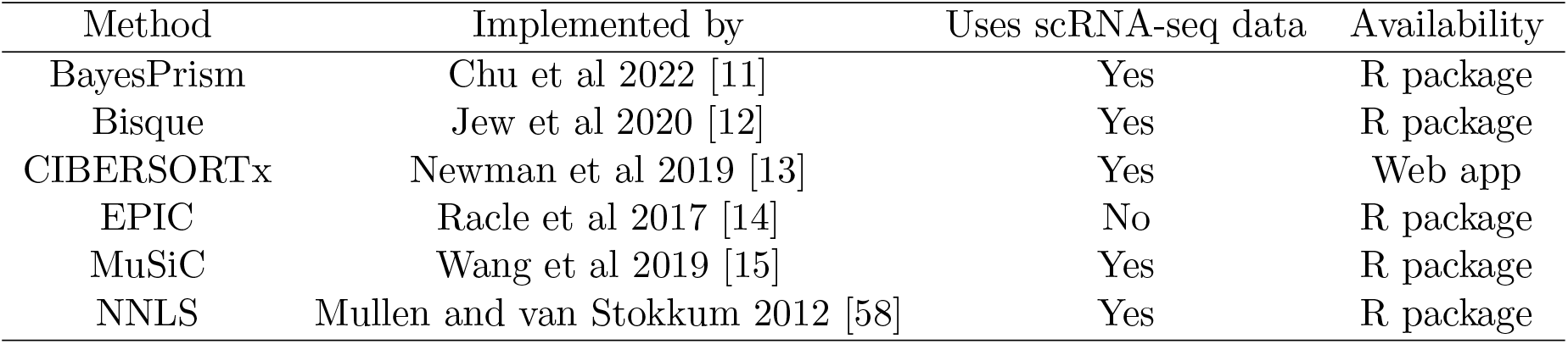
Deconvolution methods. All methods used are open source and return proportional estimates of the total composition of a tissue sample.

| Method | Implemented by | Uses scRNA-seq data | Availability |
| --- | --- | --- | --- |
| BayesPrism | Chu et al 2022 [11] | Yes | R package |
| Bisque | Jew et al 2020 [12] | Yes | R package |
| CIBERSORTx | Newman et al 2019 [13] | Yes | Web app |
| EPIC | Racle et al 2017 [14] | No | R package |
| MuSiC | Wang et al 2019 [15] | Yes | R package |
| NNLS | Mullen and van Stokkum 2012 [58] | Yes | R package |

#### Deconvolution methods have cell type bias in real and pseudo-bulk data

We generated pseudo-bulk samples using 7 of our scRNA-seq Individual samples, using cells annotated by cell type. We excluded sample 2428 due to an insufficient number of cells. We used SimBu [59] to create four datasets of 50 pseudo-bulk samples each, spanning a range of potential scenarios that a deconvolution method should be able to accurately characterize (Figure 5A). Each pseudo-bulk sample consisted of count data from 2000 single cells, sampled according to the scenario parameters. One scenario mirrored the proportions of cell types observed in the single-cell samples; we will refer to this scenario as ‘‘realistic.” Another scenario had approximately even proportions of all the cell types present in the single-cell data; we will call this scenario “even.” A scenario we will call “sparse” only included cell types believed to be common in our tumor dataset (epithelial cells, endothelial cells, fibroblasts, macrophages, and T cells), to enable us to assess how deconvolution methods handle absent cell types. One of the scenarios, called ‘weighted,” is designed to mimic our expectation that many epithelial tumors are predominated by cancer cells; in this scenario, the epithelial cell fraction was held constant at 70% with random proportions of other cell types.

**Figure 5:**
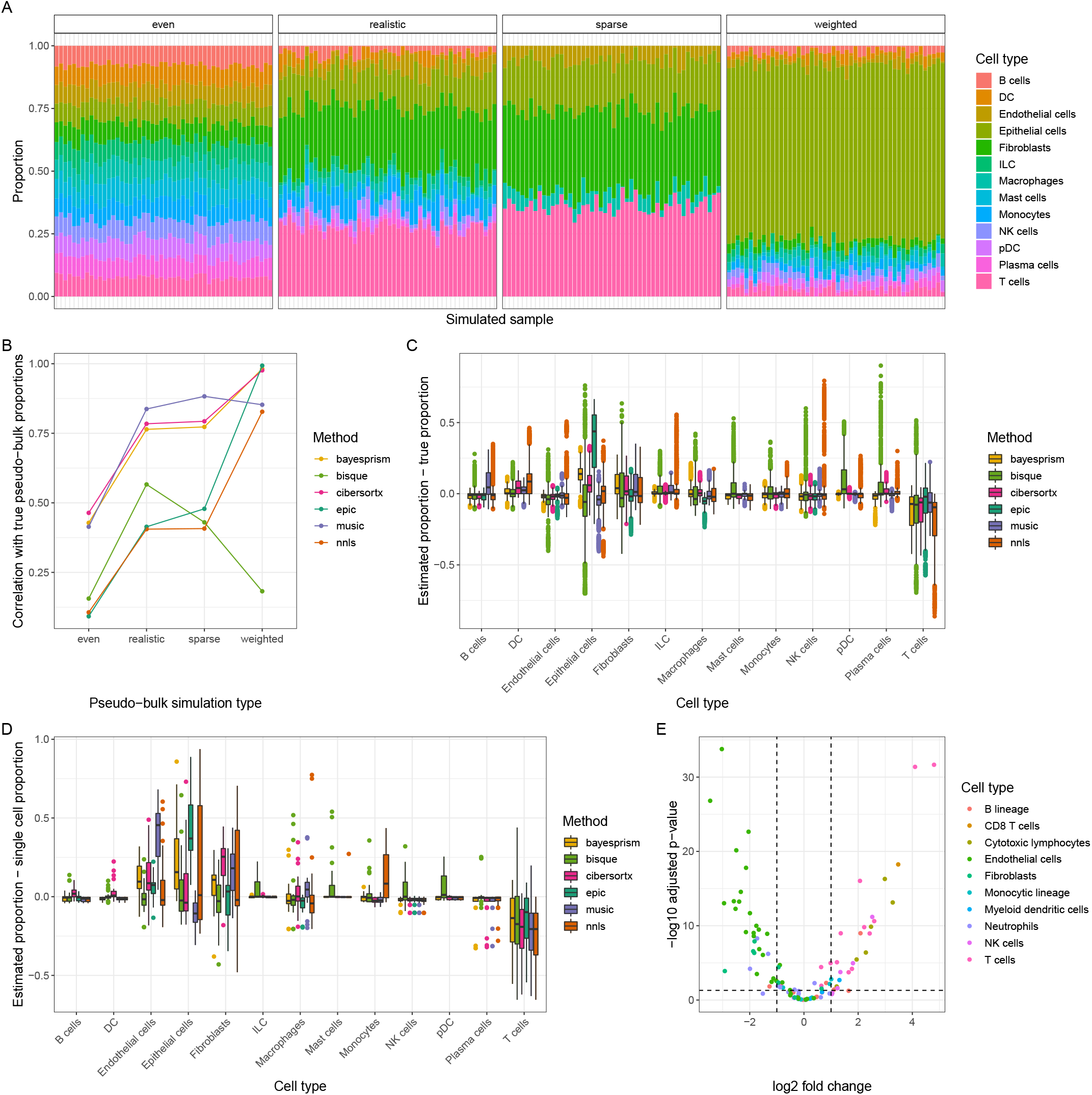
Deconvolution methods show cell type bias in pseudo-bulk and real data. A) The four different simulation scenarios to generate pseudo-bulk data from our single-cell samples, shown with a single sample. For each pseudo-bulk dataset, we pre-set proportions based on the simulation scenario and then randomly sampled cells from each cell type at those proportions. B) The average Pearson correlation value (r) between cell type proportions estimated by various deconvolution methods and the true simulated proportions, stratified by simulation type. C) The difference between deconvolution estimates for our pseudo-bulk data and their true simulated proportions, stratified by cell type. A score of 0 indicates perfect concordance between the deconvolution estimate and the true value. D) The difference between deconvolution estimates for our real data and the proportion of cells mapping to that cell type in the corresponding sample’s single-cell data. E) A volcano plot of differential expression in cell type markers between pseudo-bulk (non-simulated, aggregating all reads from all single cells in the sample) and polyA^+^ Dissociated real bulk data. Genes are selected and colored based on the signature matrix from MCPcounter [61].

We ran the deconvolution methods on all of the simulated pseudo-bulk datasets and calculated the correlation between the estimated and known proportions across samples and cell types. We found that methods differed dramatically in performance across simulation types (Figure 5B). All of the methods tested had higher correlation values in the realistic and sparse simulations than in the even simulations, the latter arguably being the least relevant to actual tissue contexts. Most of the methods trained on single-cell reference profiles performed best on the weighted simulations, whereas the methods that focus on immune cell types did poorly.

We next investigated whether some methods gave better estimates of certain cell types than others. We stratified the proportion estimates by method and cell type and subtracted the corresponding true proportion values from them (Figure 5C). Surprisingly, all the methods tested had a negative mean difference for T cells, meaning they estimated a smaller T cell fraction in the pseudo-bulk sample than was actually used. While we cannot completely explain this phenomenon, it has been reported that some kinds of T lymphocytes are underrepresented in deconvolution methods that use marker genes [60]. Perhaps the phenotypic heterogeneity observed in most T cell lineages makes it harder to identify a unifying expression profile for accurate quantification. On the other hand, most methods also overestimated the proportion of fibroblasts and epithelial cells in our data, albeit to a lesser degree than the T cells were underestimated.

Given that we had paired bulk and single-cell data, we could compare our pseudo-bulk results to the output of deconvolution on real bulk data. We ran deconvolution on each of our three bulk data types (rRNA^-^ Chunk, rRNA^-^ Dissociated, and polyA^+^ Dissociated). We used the number of cells of each type from the individual single-cell data to approximate proportions, with the assumption that deconvolution results that are closer to the proportions from the single-cell data will be closer to the unknown true proportions comprising the bulk data. (We excluded sample 2428 from this analysis because of its small number of captured single cells.) We subtracted proportions estimated from our single-cell data from the deconvolution-estimated proportions for our bulk data (Figure 5D). In this comparison, we found trends similar to those seen in the pseudo-bulk data: all methods undercount T cells, and most overcount endothelial cells, epithelial cells, and fibroblasts. These trends occur across each of the bulk data types (Figure S5A-C).

The concordance across methods led us to speculate that this may represent a true difference in cell type proportions between bulk and single-cell data. To explore this, we ran differential expression analysis on our polyA^+^ Dissociated bulk data compared to our pseudo-bulk data generated from all scRNA-seq Individual cells. Using polyA^+^ Dissociated bulk data to compare to pseudo-bulk ensured that any differences in gene expression were not an artifact of dissociation status or method of mRNA enrichment. We found a high number of differentially expressed genes, suggesting that discrepancies between bulk and single-cell data extend beyond the experimental design decisions we controlled for. We filtered differential expression results to the cell type unique markers used by MCPcounter [61]. T cell markers were all more expressed in the single-cell data than in the bulk, and the overwhelming majority of fibroblast and endothelial cell markers were more expressed in the bulk data than in the single-cell (Figure 5E). These results suggest that some step in the technical protocol post-dissociation also creates a cell type-specific bias in what cells are captured by scRNA-seq. We used microfluidic-based scRNA-seq, so loading the cells into microfluidic droplets could be differentially affected. Endothelial cells and fibroblasts are irregularly shaped and highly integrated into the extracellular matrix (ECM) and vasculature; these groups of cells may be more prone to incomplete dissociation and are strained prior to loading. This may be particularly challenging in the context of a high-grade tumor, where cancer cells establish a dense and highly disorganized ECM and vasculature compared to normal tissue. In contrast, T cells are more spherical and inherently migratory and are thus more likely to be dissociated and loaded efficiently.

Regardless of the cause of the cell type bias in scRNA-seq, its presence suggests an uncomfortable truth: bulk and single-cell RNA-seq are substantially different modalities. This challenges the use of accuracy on pseudobulk data as a gold standard for deconvolution because performing well on pseudo-bulked single-cell data does not necessarily equate to performing well on real bulk data. It also suggests that comprehensive profiles of the tumor microenvironment should include both bulk and single-cell assays to allow accurate analysis of the TME.

#### Deconvolution methods vary in robustness to technical differences

We propose an additional way to evaluate deconvolution methods: robustness of results to different experimental and protocol decisions. We posit that a method that returns consistent results for the same tissue sample, regardless of what kind of pre-sequencing processing is done and what reference profile it is given, is likely to give meaningful results across a range of real-world settings and studies. The concept of robustness has been previously employed in deconvolution with the assertion that constructing better marker gene matrices requires taking cross-microarray platform variation into account [62]. Here, we extend this concept to single-cell informed deconvolution methods.

Since our three bulk-sequenced datasets originated from the same tumors, we would expect a robust deconvolution method to return similar cell type proportions for a given tumor using each bulk dataset as input. We compared the variance in proportion estimates for each combination of sample, cell type, and method (e.g., the proportion of B cells CIBERSORTx reported for sample 2251 in rRNA^-^ Chunk vs. rRNA^-^ Dissociated vs. polyA^+^ Dissociated data) (Figure 6A). The more abundant cell types in our tumors, such as endothelial cells and epithelial cells, had naturally higher variance than the less abundant cell types, such as NK cells and plasma cells. MuSiC had the lowest variance overall, followed by BayesPrism.

**Figure 6:**
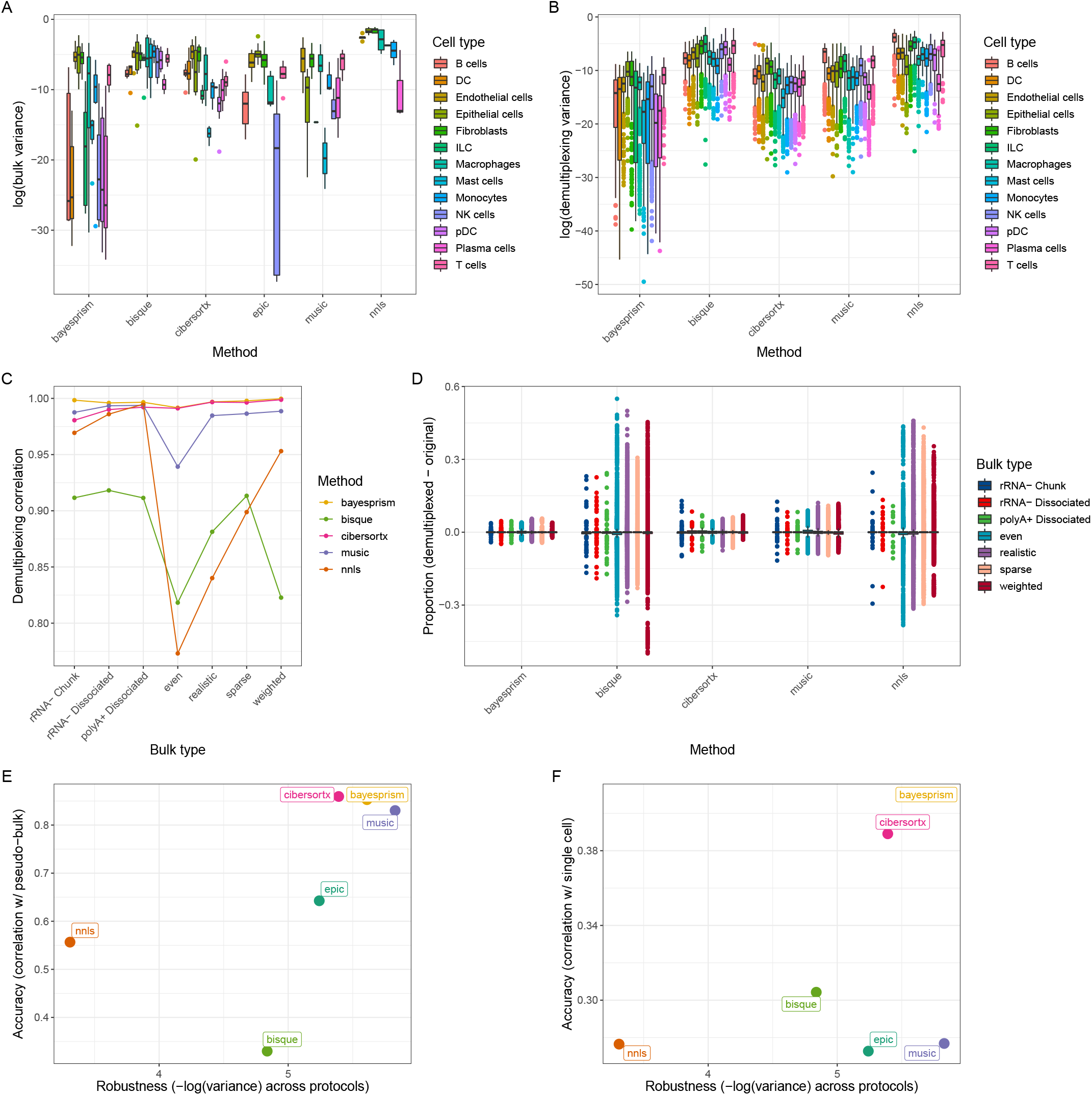
Deconvolution methods vary in robustness to changes in bulk and single-cell data. A) Variance of deconvolution results across bulk data type. For each method, we calculated the variance between the estimated proportion for a given cell type in a given sample in the rRNA^-^ Chunk, rRNA^-^ Dissociated, and polyA^+^ Dissociated data. B) Variance of deconvolution results across reference profile size. For each method, we calculated the variance between the estimated proportion of a given cell type in a given sample when using cells assigned by genetic demultiplexing (n=14,608) as a reference vs. using cells assigned by antibody-based demultiplexing with default parameters (n=7574). C) The average Pearson correlation value (r) between cell type proportions using the smaller and larger reference profile. D) The average difference between cell type proportion estimates using the smaller vs. the larger reference profile, stratified by bulk/pseudo-bulk data type. E) The final accuracy vs. robustness result for each method based on pseudo-bulk data, with variance in estimates for bulk data types and correlation between estimate and simulated proportions for pseudo-bulk data. F) Accuracy vs. robustness of each method based on true bulk data, with variance in estimates for bulk data types and correlation between real bulk estimate and real single-cell proportion.

As we have already demonstrated, changes in how the single-cell data are generated can change the cell type representation of the reference profile, which can skew deconvolution results. We used the results from our demultiplexing experiment to determine what deconvolution methods are more robust to technically-driven changes in the single-cell reference profile. We ran deconvolution on our bulk data using a reference comprising only the cells assigned by hash demultiplexing at the default 90% probability threshold. This represented 51.8% of the cells used in our original profile of cells assigned by genetic demultiplexing. Given that each cell type was still reasonably represented in the smaller single-cell dataset, we would expect a robust method to return similar deconvolution results using either reference profile. (Note that these analyses only apply to deconvolution methods that use single-cell reference profiles, so methods that use pre-selected marker genes were excluded.)

We compared the variance across the single-cell profiles in each combination of sample, cell type, method, and bulk type (e.g., the proportion of B cells CIBERSORTx reported for sample 2251’s rRNA^-^ Chunk data using the genetic demultiplexed reference profile vs. the hash demultiplexed reference profile) (Figure 6B). BayesPrism had lower average variance across most cell types. We calculated the correlation between the deconvolution results across the two reference profiles (Figure 6C). BayesPrism and CIBERSORTx had very high correlations across all bulk/pseudo-bulk types, with BayesPrism’s correlation values being nominally higher on the true bulk data. MuSiC and NNLS had comparable correlation values with the previous methods on the true bulk data types but had substantially decreased performance on the pseudo-bulk data types. Seen another way, the difference between the deconvolved cell type proportions using either reference profile was relatively small and uniform across bulk data types for BayesPrism, slightly larger but still uniform across bulk types for CIBERSORTX and MuSiC, but highly variable in Bisque and NNLS, particularly in the pseudo-bulk types (Figure 6D).

Finally, to consider robustness and accuracy simultaneously, we plotted a metric of each on an axis of a graph (Figure 6E-F) to determine if there was a tradeoff between methods, i.e., if some methods return precise-but-not-accurate results across experimental conditions and some methods are accurate under some experimental conditions but not robust. We used variance across true bulk types as the robustness axis, and for the accuracy axis we used either the correlation of pseudo-bulk proportions to true proportions (Figure 6E) or correlation of real bulk proportions to single-cell proportions (Figure 6F). We also used root mean square error (RMSE) as an accuracy metric (Figure S6A-B). BayesPrism and CIBERSORTx scored highly on both axes, and while MuSiC had a slightly higher robustness score and had high accuracy on pseudo-bulk data, it had poor accuracy on true bulk data.

## Discussion

In this study, we designed a unique experiment, profiling HGSOC tumors in multiple ways to allow for direct characterizations of how experimental design affects the deconvolution of cancer data. We introduce the metric of robustness across experimental protocols to deconvolution methods to ensure results are consistent for a single tumor independent of the technical choices made. Performing these analyses on real tumor data instead of simulated data establishes a model dataset with which future deconvolution methods can be evaluated for robustness.

We applied and evaluated six different deconvolution methods for both accuracy and robustness. We intend this to be an examination of how different commonly-used existing methods can vary in robustness and not a comprehensive benchmark. We invite researchers to use this dataset to evaluate the robustness of other existing and future methods. We have included a tutorial on GitHub for running new methods on this data (Availability of data and materials).

Our analysis focused on deconvolution methods that return absolute proportions of cell types within a sample. Other common methods return unitless scores that can be compared across samples to assess relative abundance but which do not indicate an absolute proportion of cell types in the sample. We initially applied several such methods to our data (Table S3) [61, 63–68]. However, when we attempted to assess the accuracy of these methods on our bulk data, based on their correlation with the proportions in the single-cell data, correlation values were very low (Figure S7A-G). Many of these methods focus on granular profiling of the immune compartment rather than total deconvolution, so our dataset may not be optimal for evaluating such methods.

Methods development for deconvolution is an active area of research. As such, we offer recommendations for researchers designing the next generation of deconvolution methods. One major consideration brought to light by this study is that certain cell types are present in the bulk tissue but lost from single-cell data. These cell types are thus unquantifiable by existing reference profile-based deconvolution methods. At a minimum, we recommend that future methods include a parameter to capture the proportion of “unknown cells” that lack a reference within a sample to quantify missing cell types indirectly. (This is already implemented in certain methods, such as EPIC [14]). Alternatively, a potential area for development would be a method that employs single-nucleus data (snRNA-seq) as a reference profile for deconvolution. Some cell types that are lost by dissociation can still be profiled using snRNA-seq due to the protection of the nuclear membrane [46]. A method that corrects for differences between nuclear and cytoplasmic RNA may effectively leverage an snRNA-seq reference profile to more accurately characterize all cell types present in a given tissue. Another option would be a combination of reference profile and marker gene strategies, using reference profiles for cell types that can be single-cell sequenced and cell type markers obtained from the literature or from bulk sequencing for cell types lost in single-cell sequencing.

Regardless of individual algorithmic decisions, developers of new deconvolution methods should be sure to test on real bulk and single-cell datasets that have been prepared using representative experimental protocols. As we have shown here, different design decisions each introduce biases that can affect deconvolution. Testing on only one data type renders these biases invisible. Our results show that pseudo-bulk data is an inherently limited metric and should not be used as the solitary gold standard for evaluation. Also, a recent study by Hu and Chikina confirms that the traditional way of simulating data for evaluating deconvolution does not adequately represent biological heterogeneity, and proposes new ways for better capture heterogeneity in simulation [69]. By incorporating robustness evaluations across both well-designed simulations and real datasets into their testing process, researchers can maximize the utility of their method across many future research questions.

We also have recommendations for scientists interested in designing an experiment to use deconvolution to profile the TME. For those generating novel data, pooling is an effective way to single-cell profile more tumors at a considerably reduced cost. We recommend using genetic demultiplexing to assign cells back to their sample of origin since it is independent of the efficiency of antibody loading and thus results in fewer unassigned cells with no observed bias by cell type.

Dissociation status is a key consideration when designing a bulk sequencing protocol for tumor deconvolution. Sequencing whole tumor chunks more closely approximates the tumor microenvironment than sequencing cells that have undergone dissociation due to the loss of certain cell types, but using a robust deconvolution method should allow for high performance when using either dissociated or non-dissociated bulk samples. Out of all of the methods we tested, BayesPrism had the highest combination of robustness (across bulk expression protocols and single-cell reference profile sizes) and accuracy (compared to true pseudo-bulk proportions and real single-cell data).

## Conclusion

Our results indicate that differences in data generation protocols introduce biases that alter the output of most deconvolution methods. This is true across protocols within a single data modality, such as bulk RNA sequencing of dissociated vs. non-dissociated tissue, but it is also true across different data modalities, namely bulk vs. single-cell RNA sequencing. Even when mRNA enrichment methods and dissociation status are the same, bulk and pseudo-bulk single-cell data have cell type specific abundance differences. From this, we intuit that characterizing the true cell type profile of a tissue is more complex than is deconvolving a collection of single cells that have been pseudo-bulked. Thus, accuracy on pseudo-bulk data is more of a silver standard than a gold standard. A well-performing deconvolution method will need to balance the trade-off between accuracy and robustness, being careful not to overfit to either silver standard. Out of the methods we tested, BayesPrism had the highest combination of robustness and accuracy. Development of even more robust deconvolution methods, as well as thoughtful design of experiments to generate data for deconvolution, will allow for high-quality characterizations of the TME across hundreds or thousands of samples in bulk datasets. These large sample sizes will enable a better understanding of the fundamentals of tumor biology at a population level and potentially identify opportunities for novel targeted therapeutics.

## Methods

### Experimental methods

#### Tumor processing/dissociation

Samples were collected from 8 patients with HGSOC by the University of Pennsylvania Ovarian Cancer Research Center’s Tumor BioTrust Collection (RRID: SCR_022387). All patients underwent primary debulking surgery and had not received neoadjuvant chemotherapy. A 10X enzymatic digest stock solution was made by combining a 500 mL bottle of RPMI-1640 (Gibco 61870036), 1000 mg collagenase (Millipore Sigma C9407), and 150 KU DNase type IV (Millipore Sigma D5025). Solution was sterile filtered, aliquoted, and stored at −20oC until use. Tumor samples were minced into 1mm pieces. Portions of the tumor were flash frozen. Remaining fresh tissue was put into a 1X solution of the enzymatic digest solution, diluted with RPMI-1640. Tumor tissues were dissociated overnight at room temperature. Dissociate mixture was filtered using a sterile 100*μ*m mesh filter and washed using DPBS. Red blood cells were removed using ACK Lysis Buffer. Dissociated cells were resuspended in 90% human AB serum/10% DMSO freezing media and frozen at −80C in a freezing chamber then transferred to −150C for long term storage.

#### Multiplexing

For each set of four samples, a portion of the dissociated cells was thawed and labeled with TotalSeq-B antihuman antibody-oligonucleotide conjugates from BioLegend, which are designed to label most cells via binding to both CD298 and *β*2 microglobulin. The cells were then pooled and prepared for sequencing using the 10X Genomics 3’ CellPlex Kit.

#### Single-cell sequencing

We performed scRNA-seq using the Chromium Next GEM platform from 10X Genomics. We loaded our thawed dissociated cells into emulsified droplets with Single Cell 3’ v3.1 Gel Beads using the Chromium Next GEM Chip G. We then added primers complete with a unique molecular identifier (UMI) and a poly-dT sequence to ligate with the mRNA molecules in each droplet and generate cDNA. Droplets were then broken, pooling the labeled cDNA for amplification, fragment size selection, and sequencing.

The multiplexed single-cell samples were prepared in the same way as above, but additional primers mapping to the cell surface protein feature barcodes were added alongside the other primers, allowing for specific amplification of the antibody-associated oligonucleotides.

All single-cell samples, multiplexed and individually run, were sequenced on an Illumina NovaSeq 6000 system using the S2 Reagent Kit v1.5 (100 cycles).

#### Bulk sequencing

For each sample, we bulk sequenced thawed tumor chunks; we also bulk sequenced a portion of the thawed dissociated cells in two ways: (1) Tumor chunks and one set of dissociated cells were prepared following Illumina’s TruSeq Stranded Total RNA protocol. Ribosomal RNA was depleted using the Illumina Stranded RiboZero Plus kit. Then cDNA was synthesized from the remaining RNA and enriched using PCR. (2) Another set of thawed dissociated cells were prepared according to Illumina’s TruSeq Stranded mRNA protocol. In this protocol, mRNA molecules attach to oligo-dT magnetic beads for purification before cDNA synthesis and enrichment.

All bulk samples were sequenced on an Illumina NovaSeq 6000 system using an S2 Reagent Kit v1.5 (300 cycles).

### Computational methods

#### Data processing

The single-cell data were processed using 10x Genomics’ Cell Ranger software version 6.1.2. The raw sequence files were converted to FASTQ files using the cellranger mkfastq function, which were then aligned, filtered, counted, and converted into a gene by cell matrix by the cellranger count function. These samples were aligned using a GrCh38 reference genome provided by 10x Genomics (2020-A).

The bulk data were processed using two different aligners in order to account for some deconvolution methods requiring raw read counts and others requiring transcripts per million (TPM): (1) For methods requiring raw read counts, we processed the samples using the STAR aligner version 2.7.10 [70], using an index generated from the same reference genome as the single-cell data (10x Genomics GrCh38 2020-A). We used STAR’s quantMode parameter to generate per-gene read counts for each bulk sample. (2) Since calculating transcripts per million requires consideration of transcript length, we also quantified the bulk samples using salmon version 1.9.0 [71]. We used an index generated from the GENCODE release 32 reference transcriptome (GRCh38.p13). Salmon returns per-transcript quantifications, which we combined across transcripts of a gene to get per-gene TPM values.

#### Demultiplexing

For the pooled single-cell data, we quantified and separated the cells by sample of origin using Cell Ranger version 6.1.2, specifically the function cellranger multi. In addition to the normal alignment and cell counting steps, cellranger multi also quantifies the provided cell multiplexing oligos (CMOs) and splits each cell’s read values into one of several matrices: one for each provided CMO and one for cells that weren’t able to be assigned at the given threshold. It also generates a unique BAM alignment file for the reads in each matrix and an assignment report giving the probability estimates for each barcode being assigned to each particular sample, called as a multiplet, or called as a blank droplet.

For the genetic demultiplexing, we first genotyped the STAR-aligned bulk data using the mpileup and call functions of bcftools version 1.7 [38]. To genotype the single-cell data, we used the BAM files generated by cellranger multi, concatenated into a single file. We genotyped this file using cellsnp-lite version 1.2.2 [39] with the variant calls from the bulk data as a reference for sites of heterogeneous genotypes across samples. We used vireo version 0.5.7 [40] to assign cells to a donor group based on the cellsnp-lite genotypes.

#### Single-cell processing and annotation

We used miQC to identify a sample-specific threshold using percent mitochondrial reads and library complexity (number of unique genes expressed) to filter out dead and compromised cells [72]. All cell counts reported in the paper are from after this filtering step.

We assigned cell type labels to our single-cell data, both scRNA-seq Individual and scRNA-seq Pooled, using a combination of unsupervised clustering and CellTypist [35]. For each sample and pool, we ran unsupervised clustering using the scran (version 1.24.1) and igraph (version 1.3.5) packages in R [73, 74]. Per-cluster cell type annotations were defined using marker genes, via the findMarkers function in scran. We ran CellTypist version 1.1.0 with overclustering to converge similar cells to a single cell type assignment. Cells in the pooled samples with concordant assignments based on unsupervised clustering and CellTypist were used as the default reference profile for all single cell-based deconvolution methods.

#### Bulk differential expression

Differential expression analysis was done using the DESeq2 package in R (version 1.36.0) [36]. For each analysis, we compared the eight samples from each condition (rRNA-Chunk vs. rRNA-Dissociated and rRNA-Dissociated vs. polyA+ Dissociated respectively). The sample of origin was used as a covariate to control for sample-specific effects. Principal component analysis of the bulk samples was also done using DESeq2.

#### Pseudo-bulk data generation

We used SimBu version 1.0.0 [60] as a way to efficiently sample single cells by cell type. For each scenario (even, realistic, sparse, weighted), SimBu calculated the appropriate percentage of each cell type for the custom designed scenario. For each scenario, we simulated 50 samples out of each scRNA-seq Individual sample (n=7, sample 2428 excluded). Across each simulated sample, SimBu added a random noise parameter to each cell type and then recalculated the proportions to sum to 1. It then multiplied these percentages by the desired number of cells, converted it to an integer, and then randomly sampled with replacement from the labeled single cells of that cell type. The reads from all sampled cells were then combined to form a pseudo-bulk sample. SimBu also offers a correction for different cell types having different amounts of mRNA, but we did not use this in our analysis in order to preserve the integer read counts.

#### Deconvolution

We created a snakemake pipeline [75] to run each deconvolution method on our various real and pseudo-bulk samples. We used cells from the scRNA-seq Pooled samples as a reference profile for those methods that require it. We implemented the methods that return cell type scores using the immunedeconv R package [16].

## Availability of data and materials

The dataset supporting the conclusions of this article is available in the Gene Expression Omnibus (GEO) (processed gene count tables) under accession GSE217517 and Database of Genotypes and Phenotypes (dbGaP) (raw FASTQ files) under accession phs002262.v2.p2. The code for all the analyses performed in this paper is available at https://github.com/greenelab/deconvolution_pilot under a BSD-3-Clause license.

## Ethics approval and consent to participate

All samples used in this study were obtained with patients’ written informed consent. All patients were de-identified. The protocol for tissue collection was reviewed and approved by an Institutional Review Board at the University of Pennsylvania (Protocol 702679).

## Author’s contributions

AAH: Data curation, methodology, project administration, software, visualization, writing – original draft DKO: Investigation, resources, writing – review & editing

LMW: Software, writing – review & editing

EJ: Resources, writing - review & editing

RD: Resources, writing - review & editing

JAD: Conceptualization, funding acquisition, supervision, writing – review & editing

SCH: Conceptualization, funding acquisition, supervision, writing – review & editing

CSG: Conceptualization, funding acquisition, supervision, writing – review & editing

All authors read and approved the final manuscript.

## Acknowledgements

The authors would like to thank several individuals for their assistance in this project. Thanks to Fengxiang Wang, Molly Gallagher, Zachary Berman, Jonathan Billings, and other individuals at the Center for Applied Genomics core at the University of Pennsylvania for performing sequencing of all data. Thanks to Jay Gertz (University of Utah) for advice on experimental design and common pitfalls of sequencing tumor data, as well as to Rachael Aubin for advice on running deconvolution methods. We would also like to thank Jake Crawford, Natalie Davidson, Ben Heil, and Taylor Reiter for code review.

---

**Table S1:** Number of demultiplexed cells, Batch A.

| Assignment | Hash (90% probability) | Hash (85% probability) | Hash (80% probability) | Genetic |
| --- | --- | --- | --- | --- |
| 2380 | 920 | 958 | 966 | 1010 |
| 2267 | 1119 | 1149 | 1159 | 1106 |
| 2283 | 578 | 1204 | 1737 | 2862 |
| 2293 | 1629 | 1663 | 1674 | 1720 |
| Unassigned | 2823 | 2095 | 1533 | 48 |
| Multiplets | 286 | 286 | 286 | 612 |

**Table S2:**
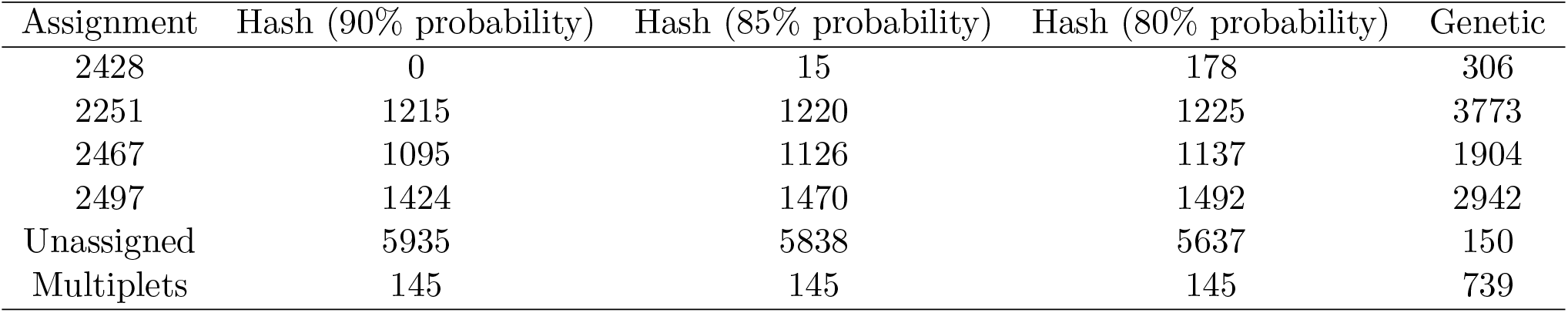
Number of demultiplexed cells, Batch B.

| Assignment | Hash (90% probability) | Hash (85% probability) | Hash (80% probability) | Genetic |
| --- | --- | --- | --- | --- |
| 2428 | 0 | 15 | 178 | 306 |
| 2251 | 1215 | 1220 | 1225 | 3773 |
| 2467 | 1095 | 1126 | 1137 | 1904 |
| 2497 | 1424 | 1470 | 1492 | 2942 |
| Unassigned | 5935 | 5838 | 5637 | 150 |
| Multiplets | 145 | 145 | 145 | 739 |

**Table S3:** Cell score deconvolution methods. All methods used are open source and return cell type scores, with a focus on immune cells.

| Method | Implemented by | Uses scRNA-seq data | Availability |
| --- | --- | --- | --- |
| ABIS | Monaco et al 2019 [61] | No | R package |
| Consensus_TME | Jiménez-Sánchez et al 2019 [62] | No | R package |
| ImmuCellAI | Miao et al 2020 [63] | No | Web app |
| MCPcounter | Becht et al 2016 [64] | No | R package |
| quanTIseq | Finotello et al 2019 [65] | No | Bash script |
| TIMER | Li et al 2020 [66] | No | R package |
| XCell | Aran et al 2017 [67] | No | R package |

**Figure S1:**
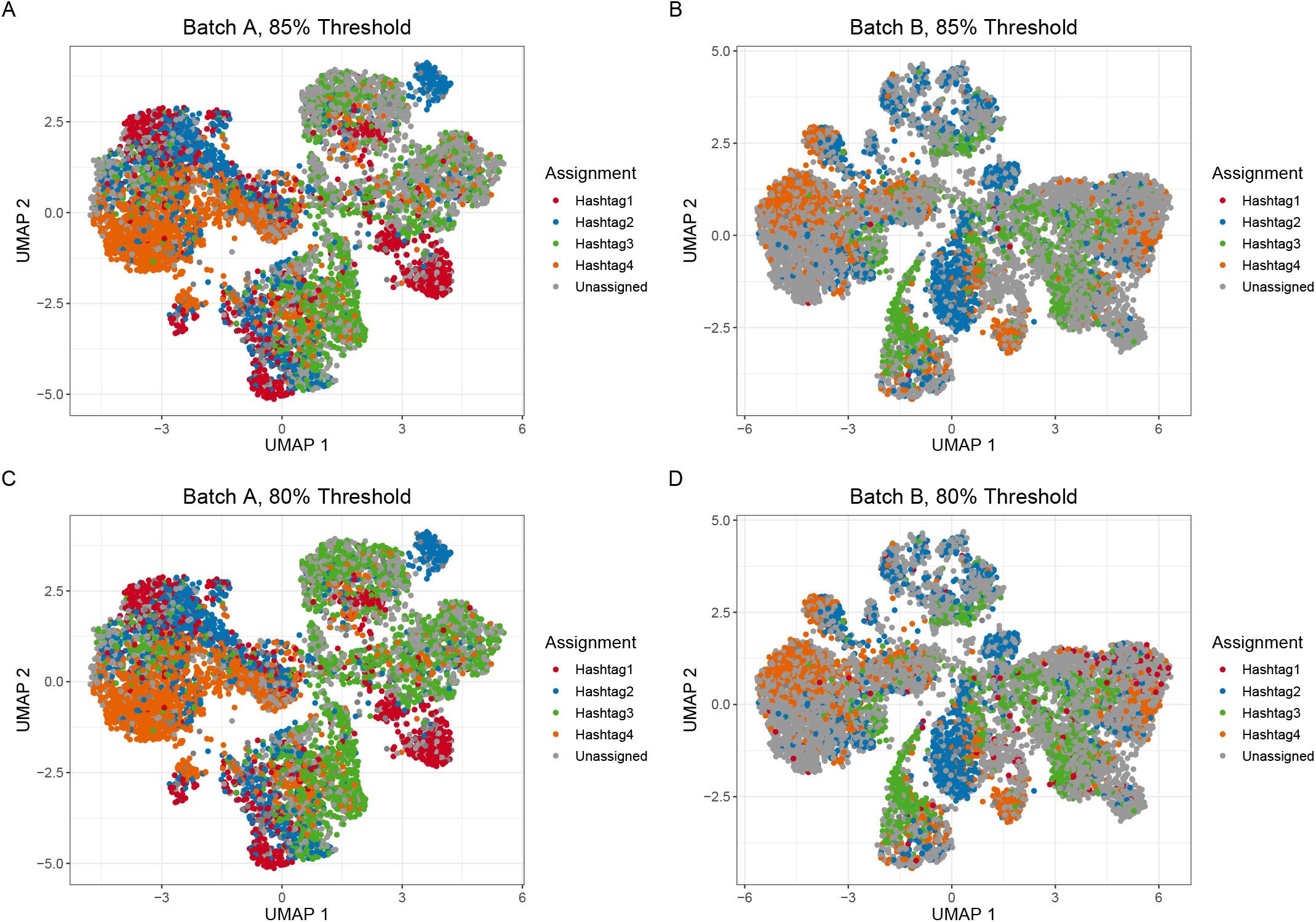
Relaxed probability thresholds for hash demultiplexing increase number of assigned cells. A) Assignments for Batch A where any cell with greater than 85% probability of originating from a sample is assigned to that sample. B) Assignments for Batch B at the 85% probability threshold. C) Assignments for Batch A at a threshold of greater than 80% probability of originating from a sample. D) Assignments for Batch B at the 80% probability threshold.

**Figure S2:**
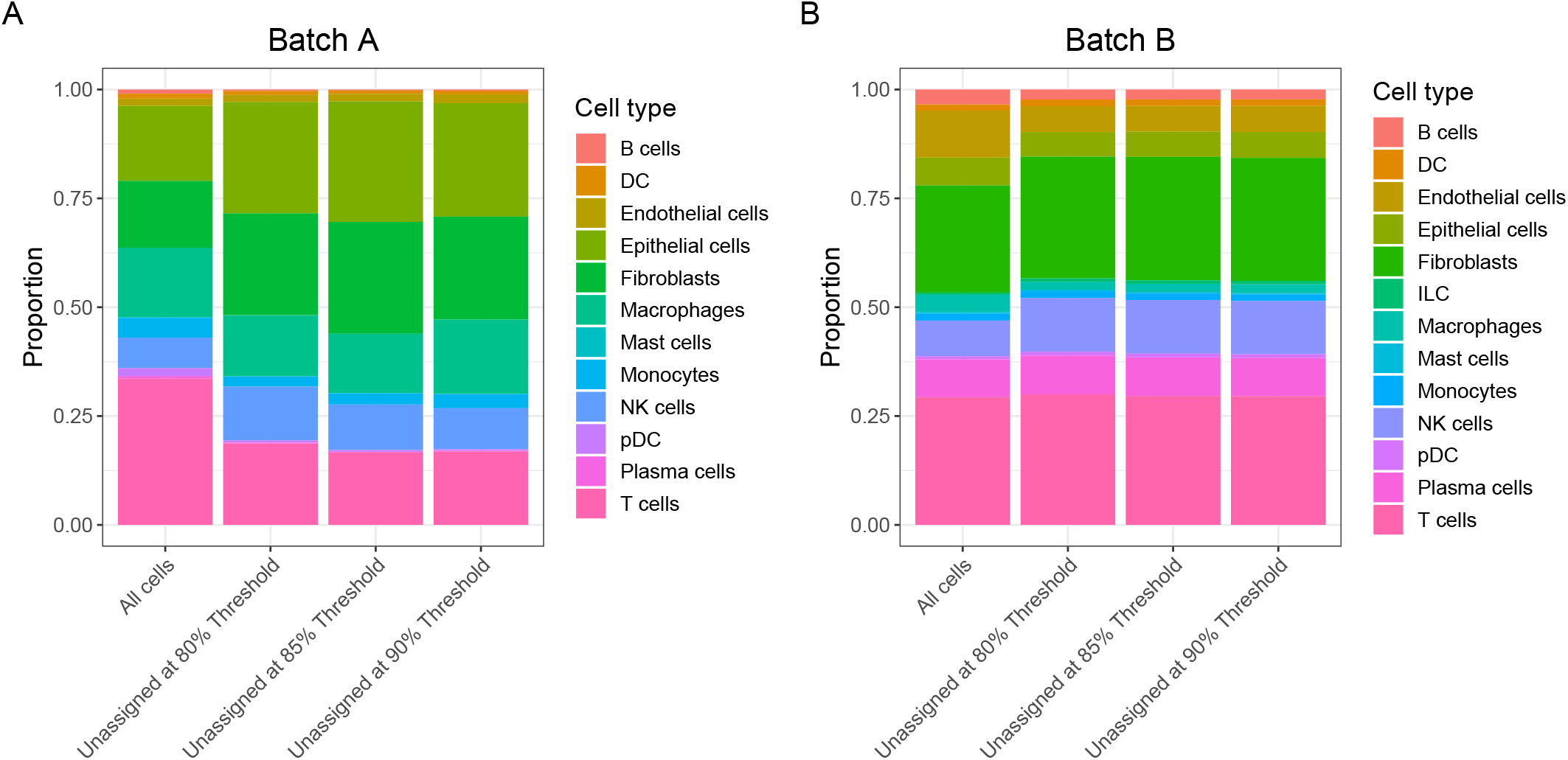
Hash demultiplexing demonstrates cell type bias. A) Proportion of cell types in Batch A across all cells and in unassigned cells at various probability thresholds. Epithelial cells and fibroblasts are proportionally greater and T cells proportionally lesser in unassigned cells than in all cells. B) Proportion of cell types in Batch B.

**Figure S3:**
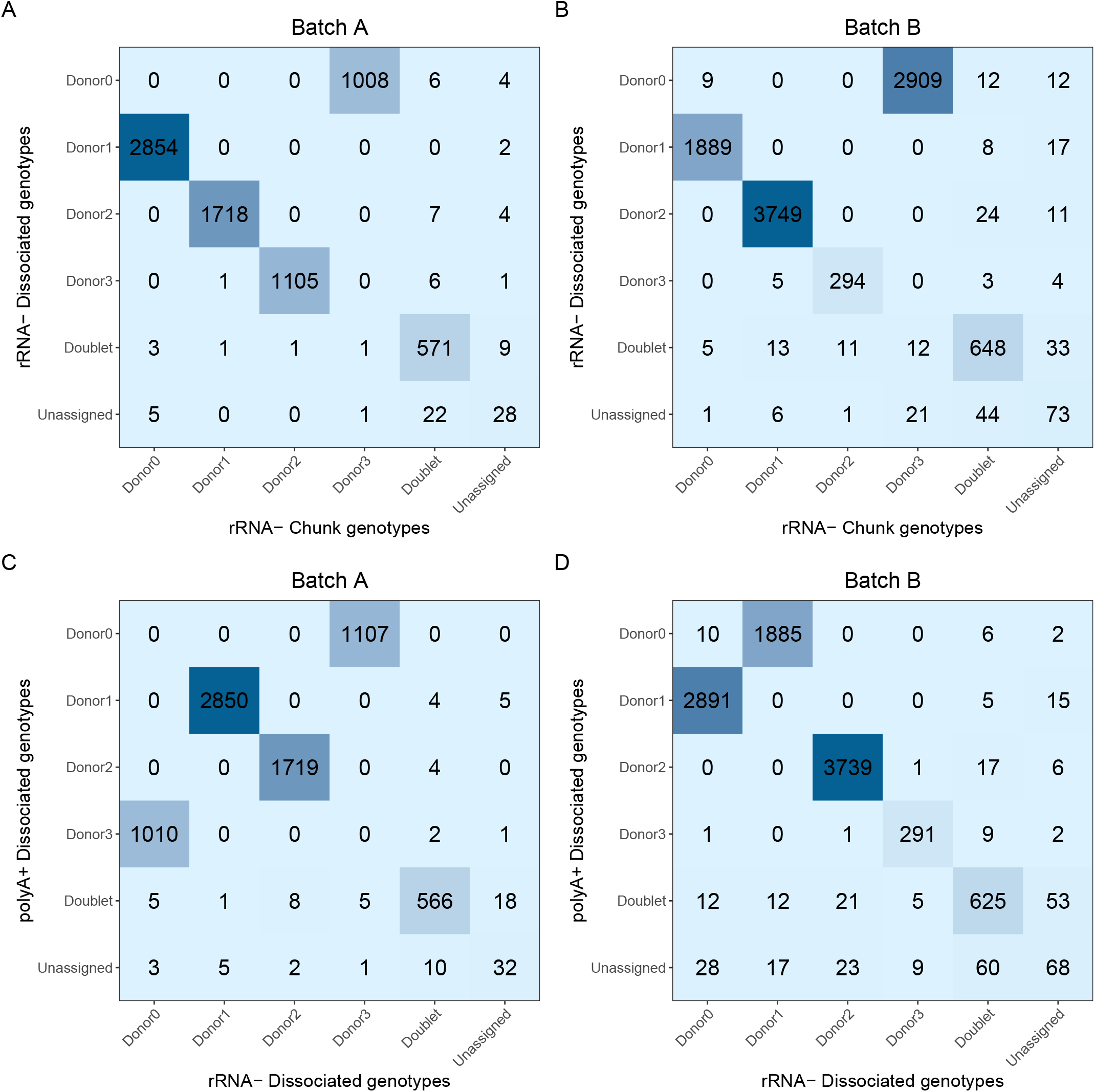
Genetic demultiplexing is concordant across source of bulk reference genotypes. A) Confusion matrix of genetic demultiplexing assignments for Batch A when using reference genotypes from rRNA^-^ Chunk samples vs rRNA^-^ Dissociated samples. B) Genetic demultiplexing assignments for Batch B using reference genotypes from rRNA^-^ Chunk samples vs rRNA^-^ Dissociated samples. C) Confusion matrix of genetic demultiplexing assignments for Batch A when using reference genotypes from rRNA^-^ Dissociated samples vs polyA^+^ Dissociated samples. D) Genetic demultiplexing assignments for Batch B using reference genotypes from rRNA^-^ Dissociated samples vs polyA^+^ Dissociated samples.

**Figure S4:**
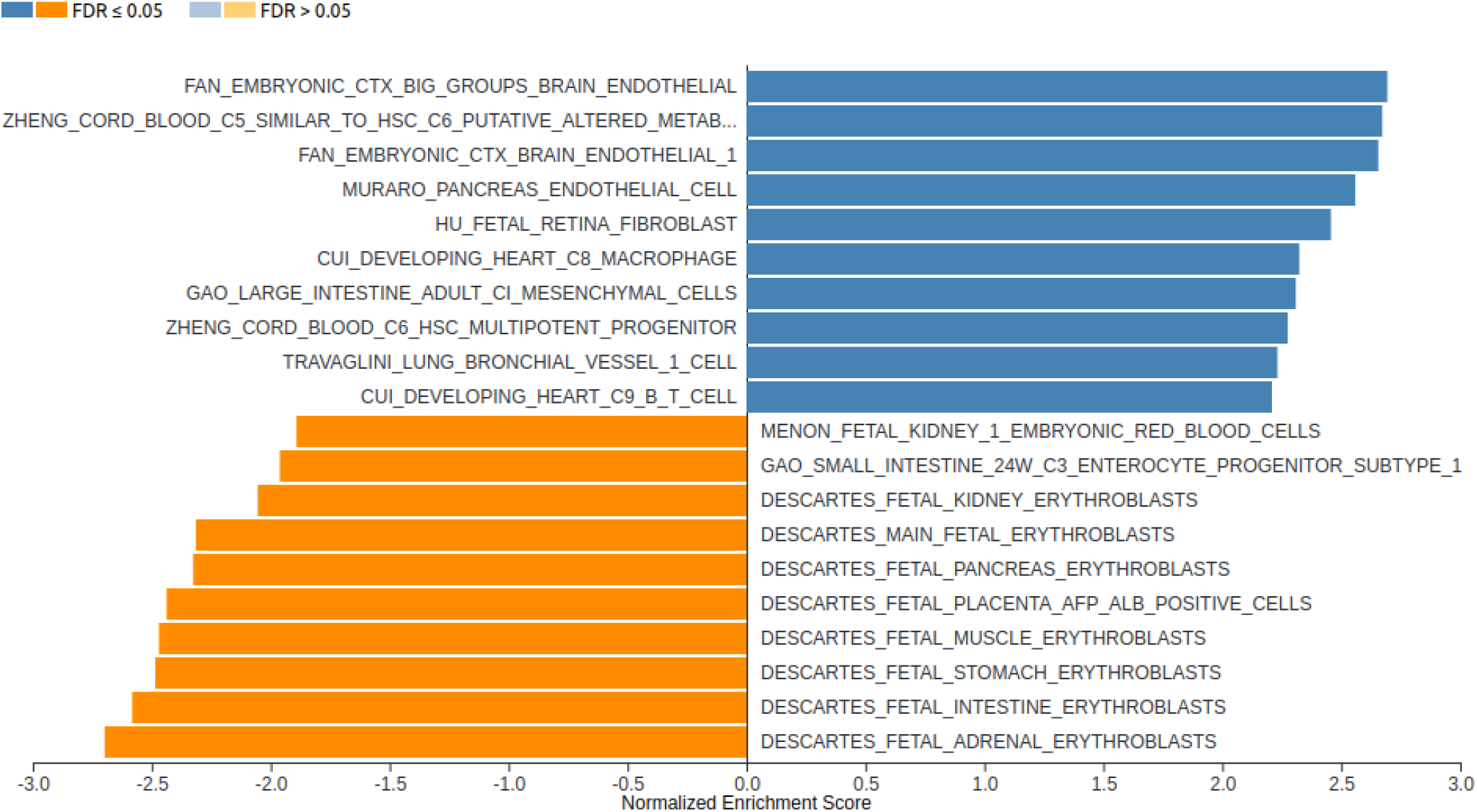
Stromal cell types are more abundant in dissociated bulk samples. Results from Gene Set Enrichment Analysis of rRNA^-^ Chunk samples vs rRNA^-^ Dissociated samples. Gene signatures associated with endothelial cells, fibroblasts, macrophages, and other immune cells (blue) are more abundant in rRNA^-^ Dissociated samples, whereas red blood cell gene signatures (orange) are more abundant in rRNA^-^ Chunk samples.

**Figure S5:**
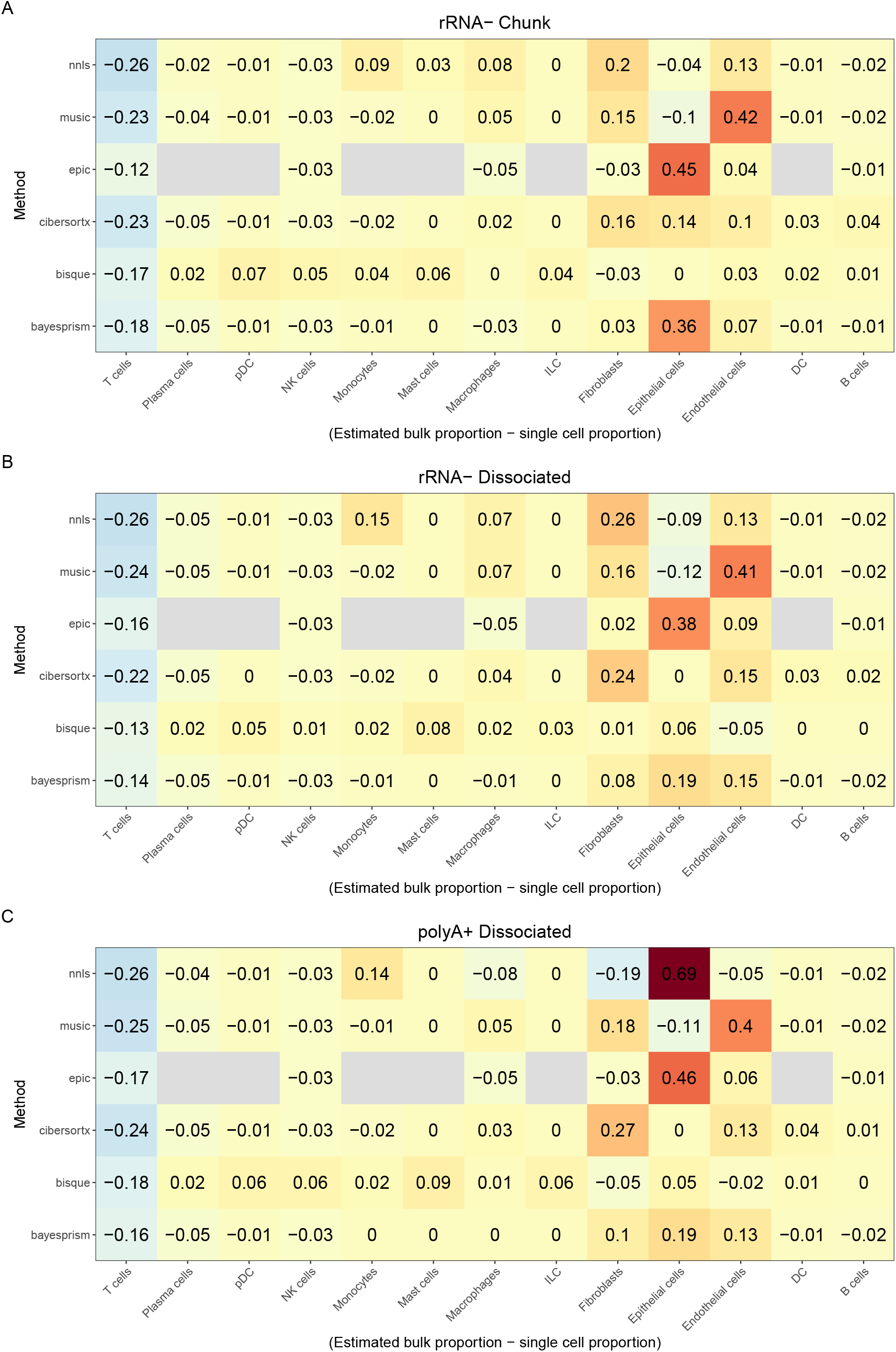
Deconvolution estimates vary based on input bulk type. A) The average difference between estimated cell type proportion in rRNA^-^ Chunk samples minus the corresponding cell type proportion in scRNA-seq Individual samples. Gray boxes represent cell types not estimated by a given method. B) The difference in cell type proportion based on rRNA^-^ Dissociated sample deconvolution estimates and scRNA-seq Individual samples. C) The difference in cell type proportion based on polyA^+^ Dissociated sample deconvolution estimates and scRNA-seq Individual samples.

**Figure S6:**
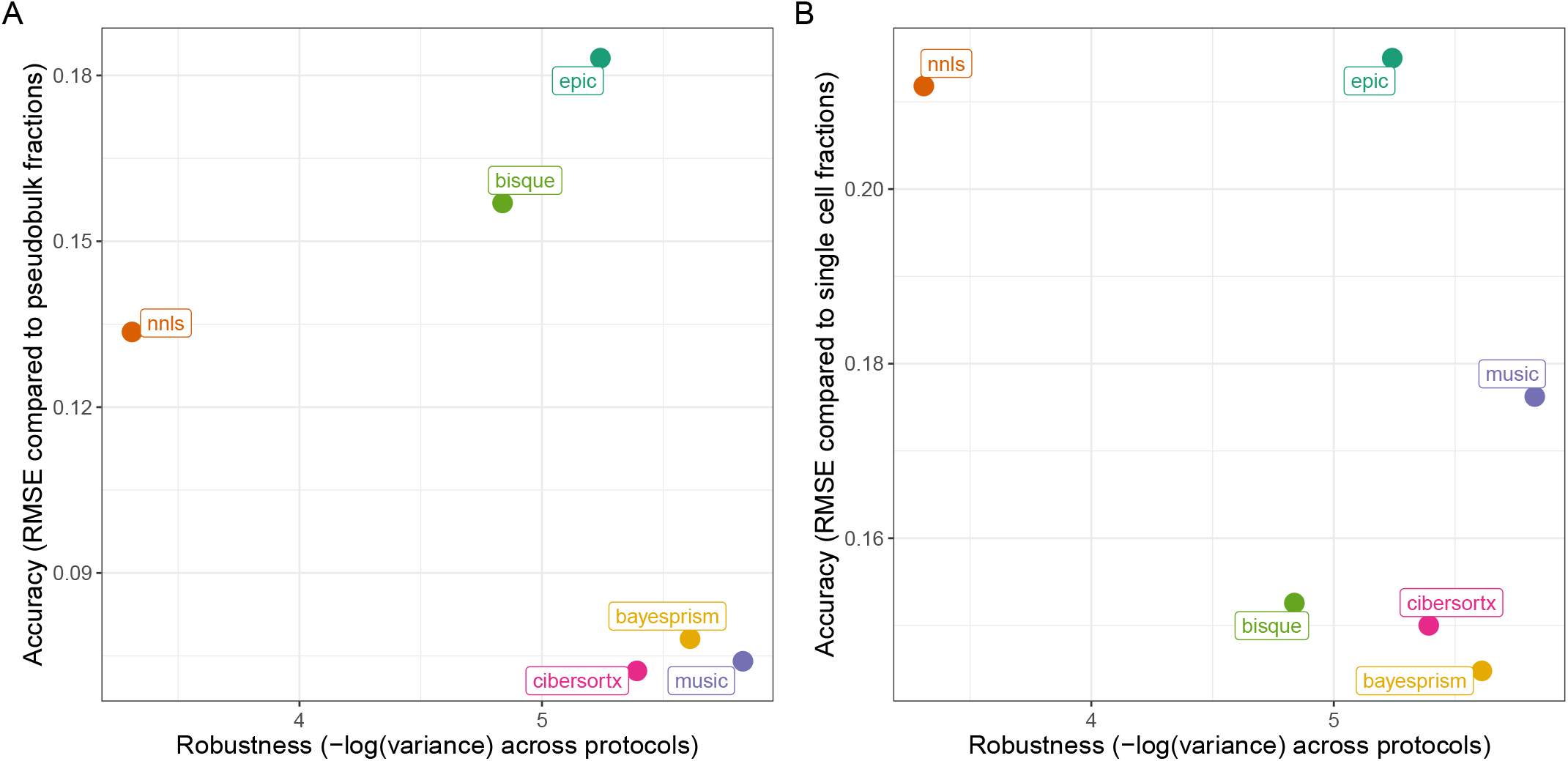
Robustness vs accuracy in deconvolution methods. A) The X axis represents the variance of estimates across the three true bulk types. The Y axis represents the root mean square error (RMSE) of pseudobulk deconvolution estimates compared to the true pseudo-bulk proportions. B) The RMSE of deconvolution proportion estimates compared to the cell type proportions in the scRNA-seq Individual samples.

**Figure S7:**
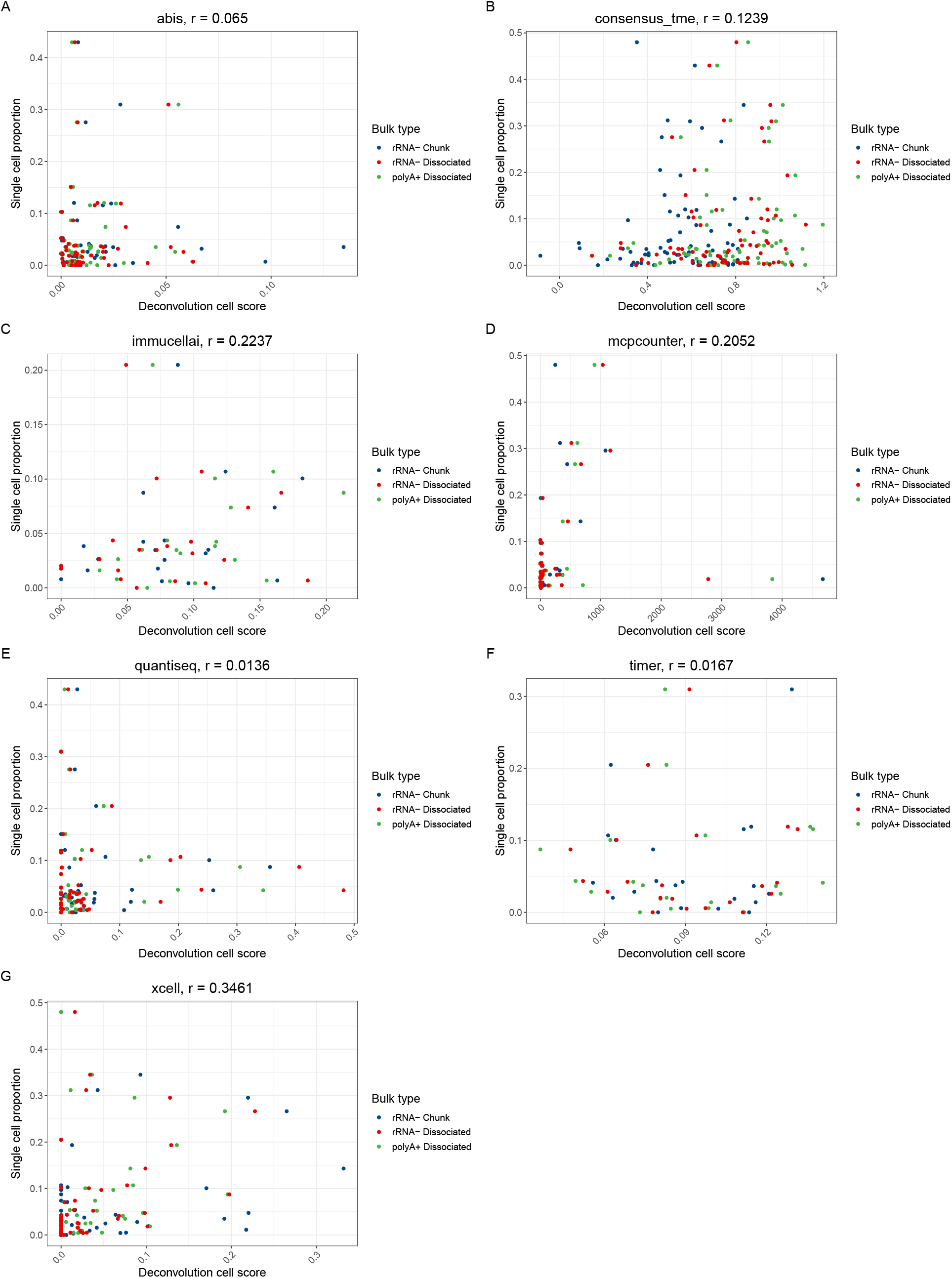
Alternate deconvolution methods that return cell type scores do not match single-cell proportions. A-G) Correlation between the cell type score returned by the deconvolution method and the corresponding proportion of cells in the scRNA-seq Individual sample. The name of the deconvolution method and the Pearson correlation (r value) is shown at the top of each panel.

## References

[1] Douglas Hanahan. Hallmarks of Cancer: New Dimensions. Cancer Discovery, 12(1):31–46, January 2022. ISSN 2159-8274, 2159-8290. doi:10.1158/2159-8290.CD-21-1059. URL https://cancerdiscovery.aacrjournals.org/content/12/1/31. Publisher: American Association for Cancer Research Section: Review.

[2] Melissa R. Junttila and Frederic J. de Sauvage. Influence of tumour micro-environment heterogeneity on therapeutic response. Nature, 501(7467):346–354, September 2013. ISSN 1476-4687. doi:10.1038/nature12626. URL http://www.nature.com/articles/nature12626. Number: 7467 Publisher: Nature Publishing Group.

[3] Paola Nisticò and Gennaro Ciliberto. Biological mechanisms linked to inflammation in cancer: Discovery of tumor microenvironment-related biomarkers and their clinical application in solid tumors. The International Journal of Biological Markers, 35(1_suppl):8–11, February 2020. ISSN 1724-6008. doi:10.1177/1724600820906155.

[4] Yi Xiao and Dihua Yu. Tumor microenvironment as a therapeutic target in cancer. Pharmacology & Therapeutics, 221:107753, May 2021. ISSN 1879-016X. doi:10.1016/j.pharmthera.2020.107753.

[5] Oliver Van Oekelen and Alessandro Laganà. Multi-Omics Profiling of the Tumor Microenvironment. Advances in Experimental Medicine and Biology, 1361:283–326, 2022. ISSN 0065-2598. doi:10.1007/978-3-030-91836-1_16.

[6] Carolyn Hutter and Jean Claude Zenklusen. The Cancer Genome Atlas: Creating Lasting Value beyond Its Data. Cell, 173(2):283–285, April 2018. ISSN 0092-8674, 1097-4172. doi:10.1016/j.cell.2018.03.042. URL https://www.cell.com/cell/abstract/S0092-8674(18)30374-X. Publisher: Elsevier.

[7] Stephanie C. Hicks, F. William Townes, Mingxiang Teng, and Rafael A. Irizarry. Missing data and technical variability in single-cell RNA-sequencing experiments. Biostatistics (Oxford, England), 19(4): 562–578, 2018. ISSN 1468-4357. doi:10.1093/biostatistics/kxx053.

[8] Jaeil Ahn, Ying Yuan, Giovanni Parmigiani, Milind B. Suraokar, Lixia Diao, Ignacio I. Wistuba, and Wenyi Wang. DeMix: deconvolution for mixed cancer transcriptomes using raw measured data. Bioinformatics, 29(15):1865–1871, August 2013. ISSN 1367-4803. doi:10.1093/bioinformatics/btt301. URL https://academic.oup.com/bioinformatics/article/29/15/1865/264697. Publisher: Oxford Academic.

[9] Catalina V Anghel, Gerald Quon, Syed Haider, Francis Nguyen, Amit G Deshwar, Quaid D Morris, and Paul C Boutros. ISOpureR: an R implementation of a computational purification algorithm of mixed tumour profiles. BMC Bioinformatics, 16, May 2015. ISSN 1471-2105. doi:10.1186/s12859-015-0597-x. URL https://www.ncbi.nlm.nih.gov/pmc/articles/PMC4429941/.

[10] Kai Kang, Caizhi Huang, Yuanyuan Li, David M. Umbach, and Leping Li. CDSeqR: fast complete deconvolution for gene expression data from bulk tissues. BMC Bioinformatics, 22(1):262, May 2021. ISSN 1471-2105. doi:10.1186/s12859-021-04186-5. URL https://doi.org/10.1186/s12859-021-04186-5.

[11] Tinyi Chu, Zhong Wang, Dana Pe’er, and Charles G. Danko. Cell type and gene expression deconvolution with BayesPrism enables Bayesian integrative analysis across bulk and single-cell RNA sequencing in oncology. Nature Cancer, 3(4):505–517, April 2022. ISSN 2662-1347. doi:10.1038/s43018-022-00356-3. URL https://www.nature.com/articles/s43018-022-00356-3. Number: 4 Publisher: Nature Publishing Group.

[12] Brandon Jew, Marcus Alvarez, Elior Rahmani, Zong Miao, Arthur Ko, Kristina M. Garske, Jae Hoon Sul, Kirsi H. Pietiläinen, Päivi Pajukanta, and Eran Halperin. Accurate estimation of cell composition in bulk expression through robust integration of single-cell information. Nature Communications, 11(1):1971, April 2020. ISSN 2041-1723. doi:10.1038/s41467-020-15816-6. URL http://www.nature.com/articles/s41467-020-15816-6. Number: 1 Publisher: Nature Publishing Group.

[13] Aaron M. Newman, Chloé B. Steen, Chih Long Liu, Andrew J. Gentles, Aadel A. Chaudhuri, Florian Scherer, Michael S. Khodadoust, Mohammad S. Esfahani, Bogdan A. Luca, David Steiner, Maximilian Diehn, and Ash A. Alizadeh. Determining cell type abundance and expression from bulk tissues with digital cytometry. Nature Biotechnology, 37(7):773, July 2019. ISSN 1546-1696. doi:10.1038/s41587-019-0114-2. URL https://www-nature-com.proxy.library.upenn.edu/articles/s41587-019-0114-2.

[14] Julien Racle, Kaat de Jonge, Petra Baumgaertner, Daniel E Speiser, and David Gfeller. Simultaneous enumeration of cancer and immune cell types from bulk tumor gene expression data. eLife, 6:e26476, November 2017. ISSN 2050-084X. doi:10.7554/eLife.26476. URL https://doi.org/10.7554/eLife.26476. Publisher: eLife Sciences Publications, Ltd.

[15] Xuran Wang, Jihwan Park, Katalin Susztak, Nancy R. Zhang, and Mingyao Li. Bulk tissue cell type deconvolution with multi-subject single-cell expression reference. Nature Communications, 10(1):380, January 2019. ISSN 2041-1723. doi:10.1038/s41467-018-08023-x. URL https://www.nature.com/articles/s41467-018-08023-x. Number: 1 Publisher: Nature Publishing Group.

[16] Gregor Sturm, Francesca Finotello, Florent Petitprez, Jitao David Zhang, Jan Baumbach, Wolf H Fridman, Markus List, and Tatsiana Aneichyk. Comprehensive evaluation of transcriptome-based cell-type quantification methods for immuno-oncology. Bioinformatics, 35(14):i436–i445, July 2019. ISSN 1367-4803. doi:10.1093/bioinformatics/btz363. URL https://www.ncbi.nlm.nih.gov/pmc/articles/PMC6612828/.

[17] Julien Racle and David Gfeller. EPIC: A Tool to Estimate the Proportions of Different Cell Types from Bulk Gene Expression Data. In Sebastian Boegel, editor, Bioinformatics for Cancer Immunotherapy: Methods and Protocols, Methods in Molecular Biology, pages 233–248. Springer US, New York, NY, 2020. ISBN 978-1-07-160327-7. doi:10.1007/978-1-0716-0327-7_17. URL https://doi.org/10.1007/978-1-0716-0327-7_17.

[18] Francisco Avila Cobos, José Alquicira-Hernandez, Joseph E. Powell, Pieter Mestdagh, and Katleen De Preter. Benchmarking of cell type deconvolution pipelines for transcriptomics data. Nature Communications, 11(1):5650, November 2020. ISSN 2041-1723. doi:10.1038/s41467-020-19015-1. URL https://www.nature.com/articles/s41467-020-19015-1. Number: 1 Publisher: Nature Publishing Group.

[19] Cole Trapnell. Defining cell types and states with single-cell genomics. Genome Research, 25(10):1491–1498, October 2015. ISSN 1088-9051, 1549-5469. doi:10.1101/gr.190595.115. URL https://genome.cshlp.org/content/25/10/1491. Company: Cold Spring Harbor Laboratory Press Distributor: Cold Spring Harbor Laboratory Press Institution: Cold Spring Harbor Laboratory Press Label: Cold Spring Harbor Laboratory Press Publisher: Cold Spring Harbor Lab.

[20] Diether Lambrechts, Els Wauters, Bram Boeckx, Sara Aibar, David Nittner, Oliver Burton, Ayse Bassez, Herbert Decaluwé, Andreas Pircher, Kathleen Van den Eynde, Birgit Weynand, Erik Verbeken, Paul De Leyn, Adrian Liston, Johan Vansteenkiste, Peter Carmeliet, Stein Aerts, and Bernard Thienpont. Phenotype molding of stromal cells in the lung tumor microenvironment. Nature Medicine, 24(8):1277–1289, August 2018. ISSN 1546-170X. doi:10.1038/s41591-018-0096-5. URL https://www.nature.com/articles/s41591-018-0096-5. Number: 8 Publisher: Nature Publishing Group.

[21] Dominic O’Neil, Heike Glowatz, and Martin Schlumpberger. Ribosomal RNA depletion for efficient use of RNA-seq capacity. Current Protocols in Molecular Biology, Chapter 4:Unit 4.19, July 2013. ISSN 1934-3647. doi:10.1002/0471142727.mb0419s103.

[22] Radmila Hrdlickova, Masoud Toloue, and Bin Tian. RNA-Seq methods for transcriptome analysis. Wiley interdisciplinary reviews. RNA, 8(1): 10.1002/wrna.1364, January 2017. ISSN 1757-7004. doi:10.1002/wrna.1364. URL https://www.ncbi.nlm.nih.gov/pmc/articles/PMC5717752/.

[23] Wei Zhao, Xiaping He, Katherine A. Hoadley, Joel S. Parker, David Neil Hayes, and Charles M. Perou. Comparison of RNA-Seq by poly (A) capture, ribosomal RNA depletion, and DNA microarray for expression profiling. BMC genomics, 15:419, June 2014. ISSN 1471-2164. doi:10.1186/1471-2164-15-419.

[24] Laura Rodriguez de la Fuente, Andrew M. K. Law, David Gallego-Ortega, and Fatima Valdes-Mora. Tumor dissociation of highly viable cell suspensions for single-cell omic analyses in mouse models of breast cancer. STAR Protocols, 2(4):100841, December 2021. ISSN 2666-1667. doi:10.1016/j.xpro.2021.100841. URL https://www.sciencedirect.com/science/article/pii/S2666166721005475.

[25] Mario L. Suvà and Itay Tirosh. Single-Cell RNA Sequencing in Cancer: Lessons Learned and Emerging Challenges. Molecular Cell, 75(1):7–12, July 2019. ISSN 1097-2765. doi:10.1016/j.molcel.2019.05.003. URL https://www.cell.com/molecular-cell/abstract/S1097-2765(19)30356-9.

[26] Francesca Finotello and Zlatko Trajanoski. Quantifying tumor-infiltrating immune cells from transcriptomics data. Cancer immunology, immunotherapy: CII, 67(7):1031–1040, 2018. ISSN 1432-0851. doi:10.1007/s00262-018-2150-z.

[27] Benjamin Izar, Itay Tirosh, Elizabeth H. Stover, Isaac Wakiro, Michael S. Cuoco, Idan Alter, Christopher Rodman, Rachel Leeson, Mei-Ju Su, Parin Shah, Marcin Iwanicki, Sarah R. Walker, Abhay Kanodia, Johannes C. Melms, Shaolin Mei, Jia-Ren Lin, Caroline B. M. Porter, Michal Slyper, Julia Waldman, Livnat Jerby-Arnon, Orr Ashenberg, Titus J. Brinker, Caitlin Mills, Meri Rogava, Sébastien Vigneau, Peter K. Sorger, Levi A. Garraway, Panagiotis A. Konstantinopoulos, Joyce F. Liu, Ursula Matulonis, Bruce E. Johnson, Orit Rozenblatt-Rosen, Asaf Rotem, and Aviv Regev. A single-cell landscape of highgrade serous ovarian cancer. Nature Medicine, pages 1–9, June 2020. ISSN 1546-170X. doi:10.1038/s41591-020-0926-0. URL http://www.nature.com/articles/s41591-020-0926-0. Publisher: Nature Publishing Group.

[28] David P. Cook and Barbara C. Vanderhyden. Ovarian cancer and the evolution of subtype classifications using transcriptional profiling. Biology of Reproduction, June 2019. doi:10.1093/biolre/ioz099. URL https://academic.oup.com/biolreprod/advance-article/doi/10.1093/biolre/ioz099/5513989.

[29] Milena Hornburg, Mélanie Desbois, Shan Lu, Yinghui Guan, Amy A. Lo, Susan Kaufman, Ashley Elrod, Alina Lotstein, Teresa M. DesRochers, Jose L. Munoz-Rodriguez, Xingwei Wang, Jennifer Giltnane, Oleg Mayba, Shannon J. Turley, Richard Bourgon, Anneleen Daemen, and Yulei Wang. Single-cell dissection of cellular components and interactions shaping the tumor immune phenotypes in ovarian cancer. Cancer Cell, 39(7):928–944.e6, July 2021. ISSN 1535-6108, 1878-3686. doi:10.1016/j.ccell.2021.04.004. URL http://www.cell.com/cancer-cell/abstract/S1535-6108(21)00212-9. Publisher: Elsevier.

[30] Neville F. Hacker and Archana Rao. Surgery for advanced epithelial ovarian cancer. Best Practice & Research Clinical Obstetrics & Gynaecology, 41:71–87, May 2017. ISSN 1521-6934. doi:10.1016/j.bpobgyn.2016.10.007. URL https://www.sciencedirect.com/science/article/pii/S1521693416301110.

[31] Michael-Antony Lisio, Lili Fu, Alicia Goyeneche, Zu-hua Gao, and Carlos Telleria. High-Grade Serous Ovarian Cancer: Basic Sciences, Clinical and Therapeutic Standpoints. International Journal of Molecular Sciences, 20(4), February 2019. ISSN 1422-0067. doi:10.3390/ijms20040952. URL https://www.ncbi.nlm.nih.gov/pmc/articles/PMC6412907/.

[32] Giovanni Ciriello, Martin L. Miller, Bülent Arman Aksoy, Yasin Senbabaoglu, Nikolaus Schultz, and Chris Sander. Emerging landscape of oncogenic signatures across human cancers. Nature Genetics, 45(10):1127–1133, October 2013. ISSN 1546-1718. doi:10.1038/ng.2762. URL https://www.nature.com/articles/ng.2762. Number: 10 Publisher: Nature Publishing Group.

[33] Peter J. Campbell, Gad Getz, Jan O. Korbel, Joshua M. Stuart, Jennifer L. Jennings, Lincoln D. Stein, Marc D. Perry, Hardeep K. Nahal-Bose, B. F. Francis Ouellette, Constance H. Li, Esther Rheinbay, G. Petur Nielsen, Dennis C. Sgroi, Chin-Lee Wu, William C. Faquin, Vikram Deshpande, Paul C. Boutros, Alexander J. Lazar, Katherine A. Hoadley, David N. Louis, L. Jonathan Dursi, Christina K. Yung, Matthew H. Bailey, Gordon Saksena, Keiran M. Raine, Ivo Buchhalter, Kortine Kleinheinz, Matthias Schlesner, Junjun Zhang, Wenyi Wang, David A. Wheeler, Li Ding, Jared T. Simpson, Brian D. O’Connor, Sergei Yakneen, Kyle Ellrott, Naoki Miyoshi, Adam P. Butler, Romina Royo, Solomon I. Shorser, Miguel Vazquez, Tobias Rausch, Grace Tiao, Sebastian M. Waszak, Bernardo Rodriguez-Martin, Suyash Shringarpure, Dai-Ying Wu, German M. Demidov, Olivier Delaneau, Shuto Hayashi, Seiya Imoto, Nina Habermann, Ayellet V. Segre, Erik Garrison, Andy Cafferkey, Eva G. Alvarez, José María Heredia-Genestar, Francesc Muyas, Oliver Drechsel, Alicia L. Bruzos, Javier Temes, Jorge Zamora, Adrian Baez-Ortega, Hyung-Lae Kim, R. Jay Mashl, Kai Ye, Anthony DiBiase, Kuan-lin Huang, Ivica Letunic, Michael D. McLellan, Steven J. Newhouse, Tal Shmaya, Sushant Kumar, David C. Wedge, Mark H. Wright, Venkata D. Yellapantula, Mark Gerstein, Ekta Khurana, Tomas Marques-Bonet, Arcadi Navarro, Carlos D. Bustamante, Reiner Siebert, Hidewaki Nakagawa, Douglas F. Easton, Stephan Ossowski, Jose M. C. Tubio, Francisco M. De La Vega, Xavier Estivill, Denis Yuen, George L. Mihaiescu, Larsson Omberg, Vincent Ferretti, Radhakrishnan Sabarinathan, Oriol Pich, Abel Gonzalez-Perez, Amaro Taylor-Weiner, Matthew W. Fittall, Jonas Demeulemeester, Maxime Tarabichi, Nicola D. Roberts, Peter Van Loo, Isidro Cortés-Ciriano, Lara Urban, Peter Park, Bin Zhu, Esa Pitkänen, Yilong Li, Natalie Saini, Leszek J. Klimczak, Joachim Weischenfeldt, Nikos Sidiropoulos, Ludmil B. Alexandrov, Raquel Rabionet, Georgia Escaramis, Mattia Bosio, Aliaksei Z. Holik, Hana Susak, Aparna Prasad, Serap Erkek, Claudia Calabrese, Benjamin Raeder, Eoghan Harrington, Simon Mayes, Daniel Turner, Sissel Juul, Steven A. Roberts, Lei Song, Roelof Koster, Lisa Mirabello, Xing Hua, Tomas J. Tanskanen, Marta Tojo, Jieming Chen, Lauri A. Aaltonen, Gunnar Rätsch, Roland F. Schwarz, Atul J. Butte, Alvis Brazma, Stephen J. Chanock, Nilanjan Chatterjee, Oliver Stegle, Olivier Harismendy, G. Steven Bova, Dmitry A. Gordenin, David Haan, Lina Sieverling, Lars Feuerbach, Don Chalmers, Yann Joly, Bartha Knoppers, Fruzsina Molnár-Gábor, Mark Phillips, Adrian Thorogood, David Townend, Mary Goldman, Nuno A. Fonseca, Qian Xiang, Brian Craft, Elena Piñeiro-Yáñez, Alfonso Muñoz, Robert Petryszak, Anja Füllgrabe, Fatima Al-Shahrour, Maria Keays, David Haussler, John Weinstein, Wolfgang Huber, Alfonso Valencia, Irene Papatheodorou, Jingchun Zhu, Yu Fan, David Torrents, Matthias Bieg, Ken Chen, Zechen Chong, Kristian Cibulskis, Roland Eils, Robert S. Fulton, Josep L. Gelpi, Santiago Gonzalez, Ivo G. Gut, Faraz Hach, Michael Heinold, Taobo Hu, Vincent Huang, Barbara Hutter, Natalie Jäger, Jongsun Jung, Yogesh Kumar, Christopher Lalansingh, Ignaty Leshchiner, Dimitri Livitz, Eric Z. Ma, Yosef E. Maruvka, Ana Milovanovic, Morten Muhlig Nielsen, Nagarajan Paramasivam, Jakob Skou Pedersen, Montserrat Puiggròs, S. Cenk Sahinalp, Iman Sarrafi, Chip Stewart, Miranda D. Stobbe, Jeremiah A. Wala, Jiayin Wang, Michael Wendl, Johannes Werner, Zhenggang Wu, Hong Xue, Takafumi N. Yamaguchi, Venkata Yellapantula, Brandi N. Davis-Dusenbery, Robert L. Grossman, Youngwook Kim, Michael C. Heinold, Jonathan Hinton, David R. Jones, Andrew Menzies, Lucy Stebbings, Julian M. Hess, Mara Rosenberg, Andrew J. Dunford, Manaswi Gupta, Marcin Imielinski, Matthew Meyerson, Rameen Beroukhim, Jüri Reimand, Priyanka Dhingra, Francesco Favero, Stefan Dentro, Jeff Wintersinger, Vasilisa Rudneva, Ji Wan Park, Eun Pyo Hong, Seong Gu Heo, André Kahles, Kjong-Van Lehmann, Cameron M. Soulette, Yuichi Shiraishi, Fenglin Liu, Yao He, Deniz Demircioğlu, Natalie R. Davidson, Liliana Greger, Siliang Li, Dongbing Liu, Stefan G. Stark, Fan Zhang, Samirkumar B. Amin, Peter Bailey, Aurélien Chateigner, Milana Frenkel-Morgenstern, Yong Hou, Matthew R. Huska, Helena Kilpinen, Fabien C. Lamaze, Chang Li, Xiaobo Li, Xinyue Li, Xingmin Liu, Maximillian G. Marin, Julia Markowski, Tannistha Nandi, Akinyemi I. Ojesina, Qiang Pan-Hammarström, Peter J. Park, Chandra Sekhar Pedamallu, Hong Su, Patrick Tan, Bin Tean Teh, Jian Wang, Heng Xiong, Chen Ye, Christina Yung, Xiuqing Zhang, Liangtao Zheng, Shida Zhu, Philip Awadalla, Chad J. Creighton, Kui Wu, Huanming Yang, Jonathan Göke, Zemin Zhang, Angela N. Brooks, Matthew W. Fittall, Iñigo Martincorena, Carlota Rubio-Perez, Malene Juul, Steven Schumacher, Ofer Shapira, David Tamborero, Loris Mularoni, Henrik Hornshøj, Jordi Deu-Pons, Ferran Muiños, Johanna Bertl, Qianyun Guo, Abel Gonzalez-Perez, Qian Xiang, and The ICGC/TCGA Pan-Cancer Analysis of Whole Genomes Consortium. Pan-cancer analysis of whole genomes. Nature, 578(7793):82–93, February 2020. ISSN 1476-4687. doi:10.1038/s41586-020-1969-6. URL http://www.nature.com/articles/s41586-020-1969-6. Number: 7793 Publisher: Nature Publishing Group.

[34] Marlon Stoeckius, Shiwei Zheng, Brian Houck-Loomis, Stephanie Hao, Bertrand Z. Yeung, William M. Mauck, Peter Smibert, and Rahul Satija. Cell Hashing with barcoded antibodies enables multiplexing and doublet detection for single cell genomics. Genome Biology, 19(1):1–12, December 2018. ISSN 1474-760X. doi:10.1186/s13059-018-1603-1. URL https://genomebiology.biomedcentral.com/articles/10.1186/s13059-018-1603-1. Number: 1 Publisher: BioMed Central.

[35] C. Domínguez Conde, C. Xu, L. B. Jarvis, D. B. Rainbow, S. B. Wells, T. Gomes, S. K. Howlett, O. Suchanek, K. Polanski, H. W. King, L. Mamanova, N. Huang, P. A. Szabo, L. Richardson, L. Bolt, E. S. Fasouli, K. T. Mahbubani, M. Prete, L. Tuck, N. Richoz, Z. K. Tuong, L. Campos, H. S. Mousa, E. J. Needham, S. Pritchard, T. Li, R. Elmentaite, J. Park, E. Rahmani, D. Chen, D. K. Menon, O. A. Bayraktar, L. K. James, K. B. Meyer, N. Yosef, M. R. Clatworthy, P. A. Sims, D. L. Farber, K. Saeb-Parsy, J. L. Jones, and S. A. Teichmann. Cross-tissue immune cell analysis reveals tissuespecific features in humans. Science, 376(6594):eabl5197, May 2022. doi:10.1126/science.abl5197. URL https://www.science.org/doi/10.1126/science.abl5197. Publisher: American Association for the Advancement of Science.

[36] Protocol -TotalSeq™-B or -C with 10x Feature Barcoding Technology. URL https://www.biolegend.com/en-us/protocols/totalseq-b-or-c-with-10x-feature-barcoding-technology.

[37] Hyun Min Kang, Meena Subramaniam, Sasha Targ, Michelle Nguyen, Lenka Maliskova, Elizabeth McCarthy, Eunice Wan, Simon Wong, Lauren Byrnes, Cristina M. Lanata, Rachel E. Gate, Sara Mostafavi, Alexander Marson, Noah Zaitlen, Lindsey A. Criswell, and Chun Jimmie Ye. Multiplexed droplet singlecell RNA-sequencing using natural genetic variation. Nature Biotechnology, 36(1):89–94, January 2018. ISSN 1546-1696. doi:10.1038/nbt.4042.

[38] Petr Danecek, James K Bonfield, Jennifer Liddle, John Marshall, Valeriu Ohan, Martin O Pollard, Andrew Whitwham, Thomas Keane, Shane A McCarthy, Robert M Davies, and Heng Li. Twelve years of SAMtools and BCFtools. GigaScience, 10(2):giab008, February 2021. ISSN 2047-217X. doi:10.1093/gigascience/giab008. URL https://doi.org/10.1093/gigascience/giab008.

[39] Xianjie Huang and Yuanhua Huang. Cellsnp-lite: an efficient tool for genotyping single cells. Bioinformatics, 37(23):4569–4571, December 2021. ISSN 1367-4803. doi:10.1093/bioinformatics/btab358. URL https://doi.org/10.1093/bioinformatics/btab358.

[40] Yuanhua Huang, Davis J. McCarthy, and Oliver Stegle. Vireo: Bayesian demultiplexing of pooled single-cell RNA-seq data without genotype reference. Genome Biology, 20(1):1–12, December 2019. ISSN 1474-760X. doi:10.1186/s13059-019-1865-2. URL https://genomebiology.biomedcentral.com/articles/10.1186/s13059-019-1865-2.

[41] Lukas M Weber, Ariel A Hippen, Peter F Hickey, Kristofer C Berrett, Jason Gertz, Jennifer Anne Doherty, Casey S Greene, and Stephanie C Hicks. Genetic demultiplexing of pooled single-cell RNA-sequencing samples in cancer facilitates effective experimental design. GigaScience, 10(9), September 2021. ISSN 2047-217X. doi:10.1093/gigascience/giab062. URL https://doi.org/10.1093/gigascience/giab062.

[42] Michael I. Love, Wolfgang Huber, and Simon Anders. Moderated estimation of fold change and dispersion for RNA-seq data with DESeq2. Genome Biology, 15(12):550, December 2014. ISSN 1474-760X. doi:10.1186/s13059-014-0550-8. URL https://doi.org/10.1186/s13059-014-0550-8.

[43] Felipe A. Vieira Braga and Ricardo J. Miragaia. Tissue Handling and Dissociation for Single-Cell RNA-Seq. In Valentina Proserpio, editor, Single Cell Methods: Sequencing and Proteomics, Methods in Molecular Biology, pages 9–21. Springer, New York, NY, 2019. ISBN 978-1-4939-9240-9. doi:10.1007/978-1-4939-9240-9_2. URL https://doi.org/10.1007/978-1-4939-9240-9_2.

[44] Xiaowei Xie, Mengyao Liu, Yawen Zhang, Bingrui Wang, Caiying Zhu, Chenchen Wang, Qing Li, Yingying Huo, Jiaojiao Guo, Changlu Xu, Linping Hu, Aiming Pang, Shihui Ma, Lina Wang, Wenbin Cao, Shulian Chen, Qiuling Li, Sudong Zhang, Xueying Zhao, Wen Zhou, Hongbo Luo, Guoguang Zheng, Erlie Jiang, Sizhou Feng, Lixiang Chen, Lihong Shi, Hui Cheng, Sha Hao, Ping Zhu, and Tao Cheng. Single-cell transcriptomic landscape of human blood cells. National Science Review, 8(3):nwaa180, August 2020. ISSN 2095-5138. doi:10.1093/nsr/nwaa180. URL https://www.ncbi.nlm.nih.gov/pmc/articles/PMC8288407/.

[45] Gabrielle J. Benitez and Kosaku Shinoda. Isolation of Adipose Tissue Nuclei for Single-Cell Genomic Applications. Journal of visualized experiments: JoVE, (160):10.3791/61230, June 2020. ISSN 1940-087X. doi:10.3791/61230. URL https://www.ncbi.nlm.nih.gov/pmc/articles/PMC7971773/.

[46] Margo P. Emont, Christopher Jacobs, Adam L. Essene, Deepti Pant, Danielle Tenen, Georgia Colleluori, Angelica Di Vincenzo, Anja M. Jørgensen, Hesam Dashti, Adam Stefek, Elizabeth McGonagle, Sophie Strobel, Samantha Laber, Saaket Agrawal, Gregory P. Westcott, Amrita Kar, Molly L. Veregge, Anton Gulko, Harini Srinivasan, Zachary Kramer, Eleanna De Filippis, Erin Merkel, Jennifer Ducie, Christopher G. Boyd, William Gourash, Anita Courcoulas, Samuel J. Lin, Bernard T. Lee, Donald Morris, Adam Tobias, Amit V. Khera, Melina Claussnitzer, Tune H. Pers, Antonio Giordano, Orr Ashenberg, Aviv Regev, Linus T. Tsai, and Evan D. Rosen. A single-cell atlas of human and mouse white adipose tissue. Nature, 603(7903):926–933, March 2022. ISSN 1476-4687. doi:10.1038/s41586-022-04518-2. URL https://www.nature.com/articles/s41586-022-04518-2. Number: 7903 Publisher: Nature Publishing Group.

[47] Kristin M. Nieman, Hilary A. Kenny, Carla V. Penicka, Andras Ladanyi, Rebecca Buell-Gutbrod, Marion R. Zillhardt, Iris L. Romero, Mark S. Carey, Gordon B. Mills, Gökhan S. Hotamisligil, S. Diane Yamada, Marcus E. Peter, Katja Gwin, and Ernst Lengyel. Adipocytes promote ovarian cancer metastasis and provide energy for rapid tumor growth. Nature Medicine, 17(11):1498–1503, November 2011. ISSN 1546-170X. doi:10.1038/nm.2492. URL https://www.nature.com/articles/nm.2492. Number: 11 Publisher: Nature Publishing Group.

[48] Arthur Liberzon, Aravind Subramanian, Reid Pinchback, Helga Thorvaldsdóttir, Pablo Tamayo, and Jill P. Mesirov. Molecular signatures database (MSigDB) 3.0. Bioinformatics, 27(12):1739–1740, June 2011. ISSN 1367-4803. doi:10.1093/bioinformatics/btr260. URL https://doi.org/10.1093/bioinformatics/btr260.

[49] Lan Dai, Keqi Song, and Wen Di. Adipocytes: active facilitators in epithelial ovarian cancer progression? Journal of Ovarian Research, 13:115, September 2020. ISSN 1757-2215. doi:10.1186/s13048-020-00718-4. URL https://www.ncbi.nlm.nih.gov/pmc/articles/PMC7513299/.

[50] Abir Mukherjee, Chun-Yi Chiang, Helen A. Daifotis, Kristin M. Nieman, Johannes F. Fahrmann, Ricardo R. Lastra, Iris L. Romero, Oliver Fiehn, and Ernst Lengyel. Adipocyte-Induced FABP4 Expression in Ovarian Cancer Cells Promotes Metastasis and Mediates Carboplatin Resistance. Cancer Research, 80(8):1748–1761, April 2020. ISSN 0008-5472, 1538-7445. doi:10.1158/0008-5472.CAN-19-1999. URL https://cancerres.aacrjournals.org/content/80/8/1748. Publisher: American Association for Cancer Research Section: Translational Science.

[51] Sven Schuierer, Walter Carbone, Judith Knehr, Virginie Petitjean, Anita Fernandez, Marc Sultan, and Guglielmo Roma. A comprehensive assessment of RNA-seq protocols for degraded and low-quantity samples. BMC Genomics, 18(1):442, June 2017. ISSN 1471-2164. doi:10.1186/s12864-017-3827-y. URL https://doi.org/10.1186/s12864-017-3827-y.

[52] William F. Marzluff, Eric J. Wagner, and Robert J. Duronio. Metabolism and regulation of canonical histone mRNAs: life without a poly(A) tail. Nature reviews. Genetics, 9(11):843–854, November 2008. ISSN 1471-0056. doi:10.1038/nrg2438. URL https://www.ncbi.nlm.nih.gov/pmc/articles/PMC2715827/.

[53] H.-L. Cao, Z.-J. Liu, P.-L. Huang, Y.-L. Yue, and J.-N. Xi. lncRNA-RMRP promotes proliferation, migration and invasion of bladder cancer via miR-206. European Review for Medical and Pharmacological Sciences, 23(3):1012–1021, February 2019. ISSN 2284-0729. doi:10.26355/eurrev_201902_16988.

[54] Yajie Chen, Qian Hao, Shanshan Wang, Mingming Cao, Yingdan Huang, Xiaoling Weng, Jieqiong Wang, Zhen Zhang, Xianghuo He, Hua Lu, and Xiang Zhou. Inactivation of the tumor suppressor p53 by long noncoding RNA RMRP. Proceedings of the National Academy of Sciences of the United States of America, 118(29):e2026813118, July 2021. ISSN 1091-6490. doi:10.1073/pnas.2026813118.

[55] Li Yang, Michael O. Duff, Brenton R. Graveley, Gordon G. Carmichael, and Ling-Ling Chen. Genomewide characterization of non-polyadenylated RNAs. Genome Biology, 12(2):R16, 2011. ISSN 1474-760X. doi:10.1186/gb-2011-12-2-r16.

[56] Simon Haile, Richard D. Corbett, Steve Bilobram, Karen Mungall, Bruno M. Grande, Heather Kirk, Pawan Pandoh, Tina MacLeod, Helen McDonald, Miruna Bala, Robin J. Coope, Richard A. Moore, Andrew J. Mungall, Yongjun Zhao, Ryan D. Morin, Steven J. Jones, and Marco A. Marra. Evaluation of protocols for rRNA depletion-based RNA sequencing of nanogram inputs of mammalian total RNA. PLOS ONE, 14(10): e0224578, October 2019. ISSN 1932-6203. doi:10.1371/journal.pone.0224578. URL https://journals.plos.org/plosone/article?id=10.1371/journal.pone.0224578. Publisher: Public Library of Science.

[57] Tomislav Ilicic, Jong Kyoung Kim, Aleksandra A. Kolodziejczyk, Frederik Otzen Bagger, Davis James McCarthy, John C. Marioni, and Sarah A. Teichmann. Classification of low quality cells from single-cell RNA-seq data. Genome Biology, 17(1):29, February 2016. ISSN 1474-760X. doi:10.1186/s13059-016-0888-1. URL https://doi.org/10.1186/s13059-016-0888-1.

[58] Katharine M. Mullen and Ivo H. M. van Stokkum. nnls: The Lawson-Hanson algorithm for non-negative least squares (NNLS), March 2012. URL https://CRAN.R-project.org/package=nnls.

[59] Alexander Dietrich, Gregor Sturm, Lorenzo Merotto, Federico Marini, Francesca Finotello, and Markus List. SimBu: bias-aware simulation of bulk RNA-seq data with variable cell-type composition. Bioinformatics, 38(Supplement_2):ii141–ii147, September 2022. ISSN 1367-4803. doi:10.1093/bioinformatics/btac499. URL https://doi.org/10.1093/bioinformatics/btac499.

[60] Marie Tosolini, Frédéric Pont, Mary Poupot, François Vergez, Marie-Laure Nicolau-Travers, David Vermijlen, Jean-Emmanuel Sarry, Francesco Dieli, and Jean-Jacques Fournié. Assessment of tumor-infiltrating TCRV9V2 lymphocyte abundance by deconvolution of human cancers microarrays. OncoImmunology, 6(3):e1284723, March 2017. ISSN null. doi:10.1080/2162402X.2017.1284723. URL https://doi.org/10.1080/2162402X.2017.1284723. Publisher: Taylor & Francis _eprint: https://doi.org/10.1080/2162402X.2017.1284723.

[61] Etienne Becht, Nicolas A. Giraldo, Laetitia Lacroix, Bénédicte Buttard, Nabila Elarouci, Florent Petitprez, Janick Selves, Pierre Laurent-Puig, Catherine Sautés-Fridman, Wolf H. Fridman, and Aurélien de Reyniès. Estimating the population abundance of tissue-infiltrating immune and stromal cell populations using gene expression. Genome Biology, 17(1):218, October 2016. ISSN 1474-760X. doi:10.1186/s13059-016-1070-5. URL https://doi.org/10.1186/s13059-016-1070-5.

[62] Francesco Vallania, Andrew Tam, Shane Lofgren, Steven Schaffert, Tej D. Azad, Erika Bongen, Winston Haynes, Meia Alsup, Michael Alonso, Mark Davis, Edgar Engleman, and Purvesh Khatri. Leveraging heterogeneity across multiple datasets increases cell-mixture deconvolution accuracy and reduces biological and technical biases. Nature Communications, 9, November 2018. ISSN 2041-1723. doi:10.1038/s41467-018-07242-6. URL https://www.ncbi.nlm.nih.gov/pmc/articles/PMC6226523/.

[63] Gianni Monaco, Bernett Lee, Weili Xu, Seri Mustafah, You Yi Hwang, Christophe Carré, Nicolas Burdin, Lucian Visan, Michele Ceccarelli, Michael Poidinger, Alfred Zippelius, João Pedro de Magalhães, and Anis Larbi. RNA-Seq Signatures Normalized by mRNA Abundance Allow Absolute Deconvolution of Human Immune Cell Types. Cell Reports, 26(6):1627–1640.e7, February 2019. ISSN 2211-1247. doi:10.1016/j.celrep.2019.01.041. URL https://www.sciencedirect.com/science/article/pii/S2211124719300592.

[64] Alejandro Jiménez-Sánchez, Oliver Cast, and Martin L. Miller. Comprehensive Benchmarking and Integration of Tumor Microenvironment Cell Estimation Methods. Cancer Research, 79(24):6238–6246, December 2019. ISSN 0008-5472. doi:10.1158/0008-5472.CAN-18-3560. URL https://doi.org/10.1158/0008-5472.CAN-18-3560.

[65] Ya-Ru Miao, Qiong Zhang, Qian Lei, Mei Luo, Gui-Yan Xie, Hongxiang Wang, and An-Yuan Guo. ImmuCellAI: A Unique Method for Comprehensive T-Cell Subsets Abundance Prediction and its Application in Cancer Immunotherapy. Advanced Science, 7(7):1902880, 2020. ISSN 2198-3844. doi:10.1002/advs.201902880. URL https://onlinelibrary.wiley.com/doi/abs/10.1002/advs.201902880. Number: 7 _eprint: https://onlinelibrary.wiley.com/doi/pdf/10.1002/advs.201902880.

[66] Francesca Finotello, Clemens Mayer, Christina Plattner, Gerhard Laschober, Dietmar Rieder, Hubert Hackl, Anne Krogsdam, Zuzana Loncova, Wilfried Posch, Doris Wilflingseder, Sieghart Sopper, Marieke Ijsselsteijn, Thomas P. Brouwer, Douglas Johnson, Yaomin Xu, Yu Wang, Melinda E. Sanders, Monica V. Estrada, Paula Ericsson-Gonzalez, Pornpimol Charoentong, Justin Balko, Noel Filipe da Cunha Carvalho de Miranda, and Zlatko Trajanoski. Molecular and pharmacological modulators of the tumor immune contexture revealed by deconvolution of RNA-seq data. Genome Medicine, 11(1):34, May 2019. ISSN 1756-994X. doi:10.1186/s13073-019-0638-6. URL https://doi.org/10.1186/s13073-019-0638-6.

[67] Bo Li, Taiwen Li, Jun S. Liu, and X. Shirley Liu. Computational Deconvolution of Tumor-Infiltrating Immune Components with Bulk Tumor Gene Expression Data. Methods in Molecular Biology (Clifton, N.J.), 2120:249–262, 2020. ISSN 1940-6029. doi:10.1007/978-1-0716-0327-7_18.

[68] Dvir Aran, Zicheng Hu, and Atul J. Butte. xCell: digitally portraying the tissue cellular heterogeneity landscape. Genome Biology, 18(1):220, November 2017. ISSN 1474-760X. doi:10.1186/s13059-017-1349-1. URL https://doi.org/10.1186/s13059-017-1349-1.

[69] Mengying Hu and Maria Chikina. Heterogeneous pseudobulk simulation enables realistic benchmarking of cell-type deconvolution methods, January 2023. URL https://www.biorxiv.org/content/10.1101/2023.01.05.522919v1. Pages: 2023.01.05.522919 Section: New Results.

[70] Alexander Dobin, Carrie A. Davis, Felix Schlesinger, Jorg Drenkow, Chris Zaleski, Sonali Jha, Philippe Batut, Mark Chaisson, and Thomas R. Gingeras. STAR: ultrafast universal RNA-seq aligner. Bioinformatics, 29(1):15–21, January 2013. ISSN 1367-4803. doi:10.1093/bioinformatics/bts635. URL https://www.ncbi.nlm.nih.gov/pmc/articles/PMC3530905/.

[71] Rob Patro, Geet Duggal, Michael I. Love, Rafael A. Irizarry, and Carl Kingsford. Salmon provides fast and bias-aware quantification of transcript expression. Nature Methods, 14(4):417–419, April 2017. ISSN 1548-7105. doi:10.1038/nmeth.4197. URL https://www.nature.com/articles/nmeth.4197. Number: 4 Publisher: Nature Publishing Group.

[72] Ariel A. Hippen, Matias M. Falco, Lukas M. Weber, Erdogan Pekcan Erkan, Kaiyang Zhang, Jennifer Anne Doherty, Anna Vähärautio, Casey S. Greene, and Stephanie C. Hicks. miQC: An adaptive probabilistic framework for quality control of single-cell RNA-sequencing data. PLoS computational biology, 17(8): e1009290, August 2021. ISSN 1553-7358. doi:10.1371/journal.pcbi.1009290.

[73] Aaron T.L. Lun, Davis J. McCarthy, and John C. Marioni. A step-by-step workflow for low-level analysis of single-cell RNA-seq data with Bioconductor. F1000Research, 5, October 2016. ISSN 2046-1402. doi:10.12688/f1000research.9501.2. URL https://www.ncbi.nlm.nih.gov/pmc/articles/PMC5112579/.

[74] Gabor Csardi and Tamas Nepusz. The Igraph Software Package for Complex Network Research. Inter-Journal, Complex Systems:1695, November 2005.

[75] Johannes Köster and Sven Rahmann. Snakemake–a scalable bioinformatics workflow engine. Bioinformatics (Oxford, England), 28(19):2520–2522, October 2012. ISSN 1367-4811. doi:10.1093/bioinformatics/bts480.

